# Shrub and Sedge Rhizosphere Communities Display Distinct Affinities Toward Exudates and Soil Organic Matter Degradation: a Quantitative Stable Isotope Probing Analysis

**DOI:** 10.64898/2025.12.02.691861

**Authors:** Sean R. Schaefer, Steven J. Blazewicz, Hannah Holland-Moritz, Caitlin Hicks Pries, Mike Allen, Jennifer Pett-Ridge, Fernando Montaño-López, A. Stuart Grandy, William Wieder, Jessica Ernakovich

## Abstract

Warming temperatures are accelerating permafrost thaw and changing tundra vegetation, where woody shrubs are displacing sedges. Shrubs, such as *Betula nana*, and sedges, such as *Eriophorum vaginatum*, exhibit distinct life strategies including unique root-associated, or rhizosphere microbial communities. As permafrost thaws it unlocks previously unavailable carbon and nutrient sources resulting in deeper roots and a translocation of rhizosphere communities. Because permafrost microbial communities contain lower diversity and biomass than rhizosphere communities, the coalescence of rhizosphere and permafrost microbial communities could alter soil organic matter (SOM) degradation rates and increase greenhouse gas emissions. To identify metabolic strategies across distinct rhizosphere and permafrost microbial communities we conducted an isotope tracing incubation experiment. We inoculated thawed permafrost with shrub and sedge rhizosphere communities while adding exudates or water daily and compared this to an uninoculated control. After 46 days, we spiked samples with ^18^O enriched water or ^13^C enriched exudates and measured isotope incorporation into microbial DNA with quantitative stable isotope probing (qSIP). Our results indicate that exudate additions had little effect on uninoculated permafrost communities but the addition of exudates and rhizosphere inoculants had a compounding effect on respiration rates. We found that soils inoculated with shrub rhizosphere communities contained a mixture of exudate and SOM degraders while soils inoculated with sedge rhizosphere communities contained mainly SOM degraders. Finally, we found that individual microbial taxa exhibited maximum growth rates in one ‘environment’, which was a combination of microbial inoculant communities and exudate addition treatments. Our results reveal that microbial niches are strongly influenced by substrate preferences and community context, and suggest that a reduction in sedges and an expansion of shrubs may provide a mechanism by which permafrost carbon losses are mitigated through corresponding shifts in microbial communities and their substrate preferences.

## 1. Introduction

Tundra regions contain over 50% of the world’s soil organic carbon stocks (Scharlemann et al., 2014). The top portion of the soil, or active layer, undergoes annual freeze-thaw cycles while the underlying permafrost remains perennially frozen (Black 1954). As warming in the Arctic reaches over four times the global average (Rantanen et al., 2022), vast permafrost carbon stocks are expected to thaw (Pörtner et al., 2019), resulting in drastic changes to tundra landscapes, including changes in hydrology, deepening of the active layer, and shifts in plant communities (Mekonnen et al., 2022, Schuur et al., 2022). Microbial activity in permafrost is limited to small brine channels (Waldrop et al., 2025). However, when permafrost thaws, the organic matter becomes vulnerable to microbial decomposition and is often accompanied by increased microbial activity and an awakening of previously dormant microbial communities in the permafrost which increases microbial respiration and greenhouse gas emissions from the soil through microbial metabolism (Burkert et al., 2019, McDonald et al., 2024). The amount of organic carbon released from permafrost thaw as greenhouse gases remains uncertain but has the potential to initiate a positive feedback loop, accelerating warming and further permafrost thaw (Schuur et al., 2015, Ernakovich et al., 2017).

Coinciding with permafrost thaw, warmer temperatures and elevated atmospheric CO_2_ levels promote higher rates of plant productivity, as indicated by a “greening” of the Arctic and shifts in vegetation communities, such as the displacement of mosses and sedges by woody shrubs in a process called *shrubification* (Sturm et al., 2001, Myers-Smith et al., 2011). The effect of shrubification on the carbon cycle remains unresolved due to the multifaceted influence of shrub expansion on ecosystem processes, including snowpack distribution and depth, soil temperatures, and microbial activity (Sturm et al., 2005). Soil warming and permafrost thaw in sedge-dominated areas, such as *Eriophorum*, can induce net soil carbon losses through increased heterotrophic respiration (Blok et al., 2018, Feng et al., 2020) which outweighs increased carbon inputs from plants (Mauritz et al., 2017). The net carbon balance in shrub-dominated areas is less clear as increased ecosystem respiration suggests a net carbon loss (Parker et al., 2015), while high C:N and slow decomposition of woody shrub biomass suggests enhanced carbon storage (Weintraub and Schimel 2005). Increased biomass and litter inputs from shrubs will alter soil organic carbon cycling (Lynch et al., 2018, Hicks et al., 2020) and heterotrophic respiration rates (Street et al., 2020) while landscape heterogeneity, including extent of permafrost thaw, soil properties, plant communities and hydrological conditions, will ultimately determine the net carbon balance of Arctic landscapes experiencing shrubification (Mekonnen et al., 2021). Rhizosphere community composition differs between shrubs, such as *Betula nana*, and sedges, such as *Eriophorum vaginatum* (Schaefer et al., 2025); these differences in composition likely extend to rhizosphere functions which affect nutrient cycling and SOM turnover. This study aims to develop a more mechanistic understanding of how shifts in root-associated microbial communities that coincide with changes in plant communities may affect SOM degradation and soil carbon dynamics following shrubification.

The primary strategies associated with rhizosphere and permafrost microbial communities are dissimilar. Rhizosphere microbes are often described as fast-growing copiotrophs associated with root exudate degradation (Lopez et al., 2023), while permafrost microbes are slower growing oligotrophs associated with SOM degradation (Ernakovich and Wallenstein 2015, Mackelprang et al., 2017). As permafrost thaws, roots extend deeper into formerly frozen, nutrient-rich soils (Blume-Werry et al., 2019) promoting the mixing, or coalescence, of rhizosphere and permafrost microbial communities (Hewitt et al., 2019). Following coalescence, emergent microbial communities tend to resemble either the most metabolically efficient taxa from both source communities or the entire source community that is more efficient (Rillig and Mansour 2017). The dominant composition and function of the emergent microbial community after coalescence, and how strongly it resembles rhizosphere or permafrost microbial communities, will have implications on carbon cycling by propagating specific metabolic associations. For instance, if a coalesced community resembles the rhizosphere community’s preference for exudates, it may mitigate greenhouse gas emissions by consuming exudates instead of SOM. By contrast, if the coalesced community more efficiently degrades SOM than exudates, emissions may be amplified as rhizosphere microbes come into contact with freshly thawed SOM. Furthermore, because plants host distinct rhizosphere communities, it is possible that outcomes of coalescence differ between plants leading to distinct perturbations in SOM cycling.

To mechanistically understand how rhizosphere communities of shrubs and sedges influence SOM degradation in thawed permafrost, we conducted a microbial community coalescence experiment using quantitative stable isotope probing (qSIP) and quantified the taxon-specific growth rates with and without artificial root exudate additions. We inoculated thawed permafrost with shrub and sedge rhizosphere slurries (*B. nana* and *E. vaginatum*) as well as sterile control inoculum. Our specific aims included: 1) measure taxon-specific growth rates for rhizosphere and permafrost microbes, 2) identify microbial strategies associated with the degradation of root exudates and/or SOM, 3) determine whether strategies of permafrost microbes change in the presence of rhizosphere microbes, and 4) determine whether the intrusion of rhizosphere communities alters SOM degradation rates. We hypothesized that the inoculation of rhizosphere communities onto permafrost would result in increased SOM decomposition and that the taxa associated with degrading exudate or SOM carbon would be consistent across exudate addition and rhizosphere inoculant treatments, indicating a fixed life history strategy regardless of microbial community.

## 2. Materials and methods

### 2.1 Permafrost and plant collection

Permafrost and plants were collected in August 2022 when active layers were near their deepest. Collections took place at the Sagwon Hills research site in the North Slope of Alaska (GPS; 69.425 -148.694). Vegetation communities at the site consisted of moist acidic tundra (Walker et al., 1994). Mean annual temperatures at Sagwon Hills are -11.7°C while the elevation is ∼214 meters (Schaefer et al., 2025). Permafrost was collected using a Snow, Ice, and Permafrost Research Establishment (SIPRE) auger. To best represent freshly thawed permafrost, samples consisted of depths between 0-10 cm below the frozen soil-permafrost boundary at the time of collection. Samples were lyophilized and homogenized before being brought to 0.68 gravimetric water content (GWC). Plant roots from *B. nana* and *E. vaginatum* were collected within 0-10 cm of the active layer surface.

### 2.2 Experimental design

The general design of the study was a 3 x 5 factorial with three rhizosphere inoculants (*B. nana, E. vaginatum,* and uninoculated control) and five isotope treatments. Embedded within isotope treatments were exudate treatments where samples either received exudate additions or water daily (Hungate et al., 2015). Samples contained ∼1 gram of thawed permafrost (∼0.6 grams dry weight mixed with 400 µL of rhizosphere slurry or water) and were incubated at 4°C for 54 days. The final eight days of the experiment represent the isotope incubation where samples were enriched in ^13^C or ^18^O isotopes, or were natural abundance controls. To track microbial community composition throughout the experiment, microbial communities were analyzed at three time points: T_0_ immediately after inoculation (n=9), T_1_ 46 days after inoculation and before the isotope was added (n=12), and T_full_ at 54 days since inoculation and 8 days after the isotope addition (n=60). We had four replicates per treatment for the isotope samples and three replicates for pre-isotope treatments for a total of 29 samples for each inoculant and 87 samples across all inoculants (Table S1).

### 2.3 Rhizosphere extraction and inoculation onto permafrost

We obtained rhizosphere communities from the sedge and shrub species (*E. vaginatum* and *B. nana*). To prepare rhizosphere slurry extracts, we shook plant roots 15 ml Falcon tubes containing ∼10 ml of 0.9% NaCl solution. We then centrifugated at 10,000 g for 15 minutes, removed top supernatant layers, and retained ∼4 ml of concentrated solution (Schaefer et al., 2025). Extracts for each plant species consisted of 6 individual roots from 3 different plants and were combined to create two representative, homogenous slurries. Batches of inoculated permafrost were made by adding 6.4 mL of rhizosphere slurries (or sterile NaCl solution for the uninoculated controls) to 32 grams of dried permafrost over. Slurries were added over the span of 10 days to ensure GWC did not exceed 0.85 and were thoroughly mixed after each addition. During this time, samples were kept at 4°C and periodically air dried (to prevent oversaturation) by placing on ice inside a laminar flow hood. Following inoculation, samples were pre-incubated at 4°C for an additional 10 days to ensure sufficient acclimation for a total of 20 days after the initial rewetting.

### 2.4 Exudate additions

Exudate solutions were created using relative molar abundances consisting of four parts glucose, two parts oxalic acid and one part alanine (Finley et al., 2018). Exudate additions occurred daily and consisted of 10 µg of C from the solution per each addition. This was done by adding 5 µL of the solution at a concentration of 2 µg/µL of C, with the exception of the first three days of the incubation where 28 µL of a 0.35 µg/µL of C solution was added. After three days, we switched to a more concentrated solution due to low evaporation from the soil. Excess water was removed weekly from the soil by brief centrifugation (∼2600 g for 10 seconds) followed by removal of the supernatant by pipetting. GWC was maintained between 0.77 and 0.87 throughout the experiment.

### 2.5 Respiration readings

CO_2_ fluxes were measured nine times over the course of the incubation using a Picarro g2201-i isotope analyzer using factory standard settings. Daily exudate additions required the jars to be opened frequently under a laminar flow and prevented anoxic conditions. CO_2_ concentrations inside the laminar flow hood were recorded at the time of the closing of the jars and used as a baseline concentration for respiration flux calculations. The increase in CO_2_ measured since the time of closing (ΔCO_2_µmol / mol air), as well as the amount of time between measurements (t) was used to calculate respiration rates (Resp). Pressure (P) was maintained at a constant 1 atm, volume (V) was a constant 0.118 L, the gas constant (R) was 0.08206 (L * atm/ mol * K) and T was a constant 277.15 K.

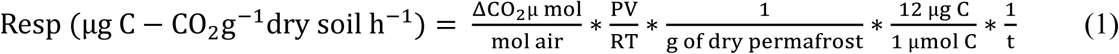

### 2.6 Calculation of soil priming

Since we did not measure SOM fluxes directly we could not measure ‘real priming’ effects, defined as the change in SOM decomposition due to plant inputs (Fontaine et al., 2003). Instead we measured ‘apparent priming’ effects, defined as the short term increases in respiration due to metabolism of exudates, SOM, and microbial turnover (Blagodatskaya and Kuzyakov 2008). Apparent priming was calculated for rhizosphere and permafrost treatments using cumulative respiration data and total carbon amendments, a technique based on the principles of root exclusion experiments (Kuzyakov 2006). Here, our treatments that did not recieve exudates served as “no-plant controls,” (“root exclusion” sensu Kuzyakov 2006) while exudate addition treatments served as our *de facto “*root inclusion.” In brief, we subtracted the moles of cumulative CO_2_ respired from our control treatments (CO_2-control_) and the moles of carbon added in exudate additions (C_exudates_) from the moles of CO_2_ respired from our exudate addition treatments (CO_2-exudate_treatment_) (equation 2). Because the exudate additions consisted of labile carbon, we assumed 60% of the carbon added was respired which is consistent with estimates using substrate specific tracers (Geyer et al., 2019). Thus, C_exudates_ was equal to the moles of carbon added via exudate additions throughout the experiment multiplied by 0.6.

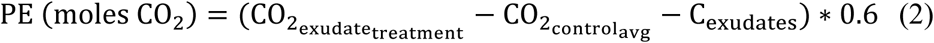

### 2.7 DNA extraction and sequencing

All soil extractions were performed using Qiagen PowerSoil Pro Kits (Qiagen, Hilden, Germany). Pre-isotope treatments (T_0_ and T_1_ samples) were extracted once while isotopically enriched samples, as well as their non-enriched counterparts (T_ful_ samples) were extracted three times each and pooled to provide enough DNA for qSIP fractionation. Approximately 10 µL of T_full_ samples were saved for amplicon sequencing (*pre*-fractionated) and the rest was used in SIP processing (*post*-fractionated). All samples were amplified for bacteria and archaea using 515f and 926r 16S primer sets (Caporaso et al., 2018) and 5.8S and ITS5FUN for fungi (Taylor et al., 2016). The thermocycler conditions for bacteria and archaea were: initial 95°C start for 2.5 minutes, followed by 35 cycles of 94°C for 45 seconds, 50°C for 1 minute and 72°C for 1.5 minute and a final annealing period of 72°C for 10 minutes. The ITS thermocycler conditions were an initial 95°C start for 2 minutes, followed by 35 cycles of 94°C for 30 seconds, 58°C for 40 seconds and 72°C for 2 minutes, and a final annealing period of 72°C for 10 minutes. We verified successful amplification on agarose gel. PCR products of 250 base pairs in length were sequenced at the Hubbard Center for Genome Studies (HCGS) at the University of New Hampshire using the Illumina NovaSeq 6000 platform.

### 2.8 Isotope enrichment

To ensure that microbial DNA contained enough isotopic enrichment for clear density gradient separation, our goal was to reach at least 50% atom percent enrichment in the H₂¹⁸O and ¹³C treatments. This was accomplished by adding 97% H_218_O water (Cambridge Isotope Laboratories, Andover, MA) or a mixture of 99 atom% ^13^C glucose (Cambridge Isotope Laboratories, Andover, MA), oxalic acid (Sigma-Aldrich, St. Louis, MO), and alanine (Sigma-Aldrich, St. Louis, MO). Prior to isotope additions, soils were dried to a GWC of 0.3; this was accomplished by centrifugation and subsequent pipetting of excess water and by placing samples on ice and circulating with moisture-free N_2_ air to increase evaporation. 180 ul of isotope solutions were added at day 46 and consisted of the following solutions: 1) natural abundance exudates and natural abundance water, 2) no exudates and natural abundance water, 3) natural abundance exudates and ^18^O-water, 4) no exudates and ^18^O-water, and 5) ^13^C exudates and natural abundance water.

To determine an appropriate incubation duration that would allow sufficient isotopic enrichment of microbial DNA for successful DNA-SIP, we first estimated how long it would take for new microbial growth to represent approximately 5–10% of the existing microbial biomass. To achieve this, we conducted a test incubation in which we measured CO₂ output from the soil over one week. By quantifying soil respiration rates during this preincubation, we were able to estimate the amount of microbial growth that occurred, assuming a relatively low carbon use efficiency (CUE) of 20%. This approach allowed us to predict the extent of isotope incorporation into newly synthesized microbial DNA during the planned SIP experiment. Specifically, we estimated initial DNA concentrations by pipetting 1.5 µL of DNA extracts onto a DeNovix DS-11+ Spectrophotometer (DeNovix, Wilmington, DE), and multiplied this value by 38% to estimate the amount of carbon in DNA (ng C). Next, using *in situ* respiration rates we calculated the amount of C respired over a period of 3 and 7 days based on minimum, median, and maximum respiration rates observed (0.15, 0.5 and 1.1 µg C * gram of dry soil^-1^ * hour^-1^). We then multiplied the amount of carbon respired by 20% carbon use efficiency and divided that by the amount of carbon in starting DNA to estimate the amount of new biomass.

### 2.9 Sequence data processing

Average sequence depths for *pre*-fractionated (T_0_ and T_1_) bacteria, and fungi were 38,838 and 61,755 respectively and ranged between 4,331 – 149,441 reads for bacteria and 2,018 – 295,824 reads for fungi. Average raw sequencing depths for *post-*fractionated (T_full_) bacteria, and fungi were 30,208 and 28,447 respectively and ranged between 2 – 980,743 reads for bacteria and 1 – 390,380 reads for fungi. It is worth noting that we did detect some archael sequences in 16S primers, however their abundance was sparse and are combined and referred to with bacteria throughout the manuscript. All sequences were processed, filtered, and trimmed using DADA2 v.1.20 (Callahan et al., 2016). Minor modifications to the pipeline were made to optimize for Novaseq sequencing platforms (Teixeira et al., 2024); this included removing any reads with 20 or more polyG’s and implementing an error rate learning step that allowed for more accurate estimation of error rates from binned quality scores, an artifact of Novaseq sequencing platforms. All primers and sequences containing 20 or more polyG’s were removed with Cutadapt v.3.5. 16S sequences were then filtered to remove low-quality reads (truncLen=c(230,230), truncQ=2, maxN=0, maxEE=c(2,2),rm.phix=TRUE)). ITS sequences were filtered but not trimmed due to the variable lengths of the PCR product (maxN = 0, maxEE = c(2, 2), truncQ = 2, minLen = 50, rm.phix=TRUE, compress=TRUE, verbose=TRUE, multithread=TRUE)). Instead, we used ITSxpress to identify and extract ITS regions prior to denoising (Rivers et al., 2018). Then, 16S and ITS sequences were denoised using DADA2 software. Taxonomy was assigned from amplicon sequence variants (ASV’s) using Naïve Bayes classifiers. The databases used were SILVA v.138.1 for bacteria and UNITE v.8.3 for fungi. In total we found 34,331 bacterial and 2,278 fungal ASVs for pre-fractionated samples and 42,511 and 13,313 for post-fractionated. Post-DADA2 processing and before subsequent analyses, we removed samples with less than 4,000 reads for bacteria and less than 2,000 for fungi, chloroplast and mitochondrial DNA sequences, and normalized abundances to median sequencing depth, all of which was done in R using the phyloseq package (v1.42.0; McMurdie and Holmes 2013). PCR and DNA extraction blanks were included in DNA sequencing to monitor background contamination. Low levels of bacterial and fungal DNA were detected in T_1_ blanks and were removed from the analysis after confirming their near zero species richness and distinct community composition from samples through ordination analysis.

### 2.10 DNA Fractionation and SIP processing

All post-incubation SIP processing was done at Lawrence Livermore National Laboratory (LLNL) using their automated High Throughput Stable Isotope Probing (HT-SIP) pipeline (Nuccio et al., 2022). In brief, DNA samples were placed into ultracentrifuge tubes containing 4.5 mL of CsCl (1.885 g*mL^-1^), 1 mL gradient buffer (0.1 mol/L Tris, 0.1 mol/L KCl, and 1 mmol/L EDTA), 5 μL gradient buffer with 0.1% Tween, and between 0.5 to 2 μg of DNA suspended in 50 μL of TE buffer (10mM Tris-HCl containing 1mM EDTA) for a final density of 1.725 - 1.730 g*mL^-1^. Samples were then spun on an ultracentrifuge for 108-136 h at 176,284 x *g* at 20 °C in a Beckman Coulter Optima XE-90 ultracentrifuge using a VTi65.2 rotor and fractionated into 22 fractions using an Agilent Technologies 1260 Isocratic Pump and 1260 Fraction Collector. During fractionation, CsCl was displaced with sterile water pumped at 0.25 mL*min^-1^ resulting in ⁓250 µL per fraction and fractionated samples were measured for density using a Reichart AR200 digital refractometer. The DNA contained in each fraction was then purified and concentrated using glycogen/ PEG precipitation followed by an ethanol wash (Dunford and Neufeld 2010). A Hamilton Microlab STAR liquid handling robot was used to automate this process where 500 uL PEG solution (30% PEG 6000, 1.6 M NaCl) and 35 µL of 1:5 diluted Glycoblue (Invitrogen, Carlsbad, CA) were added to each fraction and incubated overnight at room temperature. PEG was removed by spinning samples at 4198 RCF for 5 h at 10°C and 950 µL 70% ethanol was added to each fraction. Ethanol was then removed, and DNA was resuspended in 40 µL of 1x Tris-EDTA (pH 7.5) before DNA concentrations were quantified using a PicoGreen fluorescence assay (Invitrogen, Carlsbad, CA). Fractions were then analyzed to identify the minimum and maximum CsCl densities containing sufficient DNA (0.2 ng/µL or above). The resulting endpoints were between 1.689 and 1.748 g/mL of CsCl and resulted in 762 fractionated samples with an average of 10 fractions per soil sample.

### 2.11 Quantitative PCR

Quantitative PCR was conducted using the same primers as sequencing minus the adapters necessary for Illumina sequencing. All samples were run in triplicate in 10 µL reactions. The bacterial runs contained 5 µL of SsoAdvanced Universal SYBR Green Supermix (Bio-Rad Laboratories, Hercules, CA), 0.1 µL of forward and reverse primers (100 ng/µL; Invitrogen, Carlsbad, CA), 3.9 µL of PCR grade water (Teknova, Hollister, CA) and 1 µL of DNA. Fungal runs contained 5 µL of SsoAdvanced Univ Probes Supermix (Bio-Rad Laboratories, Hercules, CA), 0.05 µL of custom ITS TaqMan MGB Probe (0.02 umol/µL; Applied Biosystem, Woburn, MA), 2 µL of forward and reverse primers (10 ng/µL, Invitrogen, Carlsbad, CA), 1.95 µL of PCR grade water (Teknova, Hollister, CA) and 1 µL of DNA. Bacterial amplicon standard curves were created using a 465 bp DNA fragment amplified from *Salmonella enteritidis* with 341F/806R 16S rRNA primers (Klindworth et al., 2013 Apprill et al. 2015). The insert was cloned into a PCR 2.1 TOPO vector (Invitrogen, Carlsbad, CA). The linearized 4365 bp plasmid was diluted serially 10-fold and standard curves were made up of 1.7 x 10^2^ to 2.4 x 10^8^ plasmid copies per reaction. Fungal standard curves were created using ITS4 and 5.8s-FUN primers and ITS TaqMan MGB Probe sequences which were inserted into a pTWIST Amp High Copy vector (Twist Bioscience, San Francisco, CA). The primer and probe sequences were placed at nucleotide positions mimicking their relative positions within the *Serendipita vermifera* genome and were flanked by random DNA sequence for a total insert size of 943 bp. The constructed plasmid was linearized and diluted serially 10-fold and standard curves were made up of 4.4 x 10^1^ to 8.8 x 10^8^ plasmid copies per reaction. Annealing temperatures were tested at intervals of 0.5°C between 51°C and 60°C to determine ideal annealing temperatures; target regions were confirmed using a melt curve. Thermocycler conditions for bacterial qPCR consisted of an initial start of 98 °C for 3 minutes, followed by a cycle of 98 °C for 15 seconds and 55 °C for 30 seconds, and was repeated for 40 rounds. Thermocycler conditions for fungal qPCR consisted of an initial start at 98 °C for 3 minutes, followed by a cycle of 98 °C for 15 seconds, 56.7 °C for 30 seconds and 72 °C for 45 seconds, and was repeated for 40 rounds. Average PCR efficiency was 97.8% for bacteria and 89.0% for fungi. Average slopes were -3.38 and -3.63 for bacteria and fungi respectively and all standard curves contained an R^2^ value ≥ 0.99.

### 2.12 Quantitative Stable Isotope Probing (qSIP) calculations

All qSIP calculations were performed using the R package qSIP2 (v.0.18.4.9001; Kimbrel 2024). Taxon-specific shifts in density caused by the incorporation of ^18^O and ^13^C isotopes into microbial DNA, or enriched atom fraction (EAF) values, were calculated using qSIP procedures described previously (Hungate et al., 2015, Koch et al., 2018, Blazewicz et al., 2020). In brief, EAF values correspond to a difference in taxon-specific densities between DNA that is isotopically enriched, compared to DNA of natural abundance isotopes. The difference in taxon-specific densities is directly proportional to the assimilation of isotope into that individual taxon’s newly synthesized DNA. Microbes with a detectable density shift were considered active while undetected microbes are considered dormant or inactive.

We estimated the number of 16S and ITS rRNA gene copies per taxon in each density fraction by multiplying the relative abundances of each taxon (derived from DNA sequencing of fractionated samples) by the total number of 16S or ITS gene copies obtained from qPCR. Because we were interested in rare, but not singularly occurring taxa, we chose to only include taxa that occurred in at least 2 (of 12) replicates and at least 2 fractions. All EAF values and 95% confidence intervals were calculated using bootstrapping of 1000 iterations (Hungate et al., 2015).

Since DNA replication is directly proportional to cell division, we used newly synthesized DNA as a proxy for cell growth. We assumed a constant proportion of heavy oxygen, derived from water, is integrated into newly synthesized DNA (propO= 0.6). Likewise, we assumed a constant proportion of heavy carbon, derived from synthetic exudates, is integrated into newly synthesized DNA (propO=1.0). These assumptions were based on experimental evidence and mathematical constraints (Blazewicz and Schwartz 2011, Hungate et al., 2015, Koch et al., 2018).

The qSIP model assumes that all newly synthesized DNA will be isotopically labeled; thus, increases in 16S or ITS marker genes were used for per capita growth rates. Absolute population growth rates were used to compare ^13^C and ^18^O isotope treatments (Koch et al., 2018, Blazewicz et al., 2020); however, it is worth noting that ^13^C growth rates are proportionally related and are not truly absolute population growth rates. We estimated growth rates by calculating the change in 16S and ITS gene copies per gram of soil over the final eight days of the incubation (equation 3). Using the molecular weights of our isotopically labeled and unlabeled DNA we determined the number of gene copies per gram of dry soil.

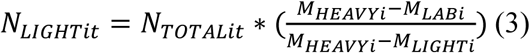

Where *N_LIGHTit_* is the number of unlabeled 16S or ITS genes, and *N_TOTALit_* is the number of labeled plus unlabeled 16S or ITS genes of taxon *i* at the end of the incubation (*t*). *M_HEAVYi_* and *M_LIGHTi_* are the molecular weights of ^13^C or ^18^O labeled DNA, and unlabeled DNA of taxon *i,* respectively. *M_LABi_* is the average molecular weight of labeled and unlabeled DNA at the end of the incubation. From these, we were able to calculate the growth rates of taxon *i* (equation 4).

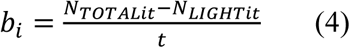

Where *b_i_* is the growth rate of taxon *i* over the final eight days of the incubation, *t*.

### 2.13 Substrate preferences and strategy designations

Specific comparisons of growth rates between the treatments reveal information about what type of substrate a microbe prefers within an environment (Fig. 1, Table S2). We used three assumptions to guide our designations: 1) microbial turnover and necromass recycling was minimal following the eight days after isotope solutions were added into the soils, 2) the incorporation of ^13^C into DNA represents exudate consumption rates that are proportionally related to growth rates but are not total gross population growth rates, and 3) incorporation of ^18^O-water into DNA is representative of total gross population growth rates (Blazewicz and Schwartz 2011).

**Fig. 1.**
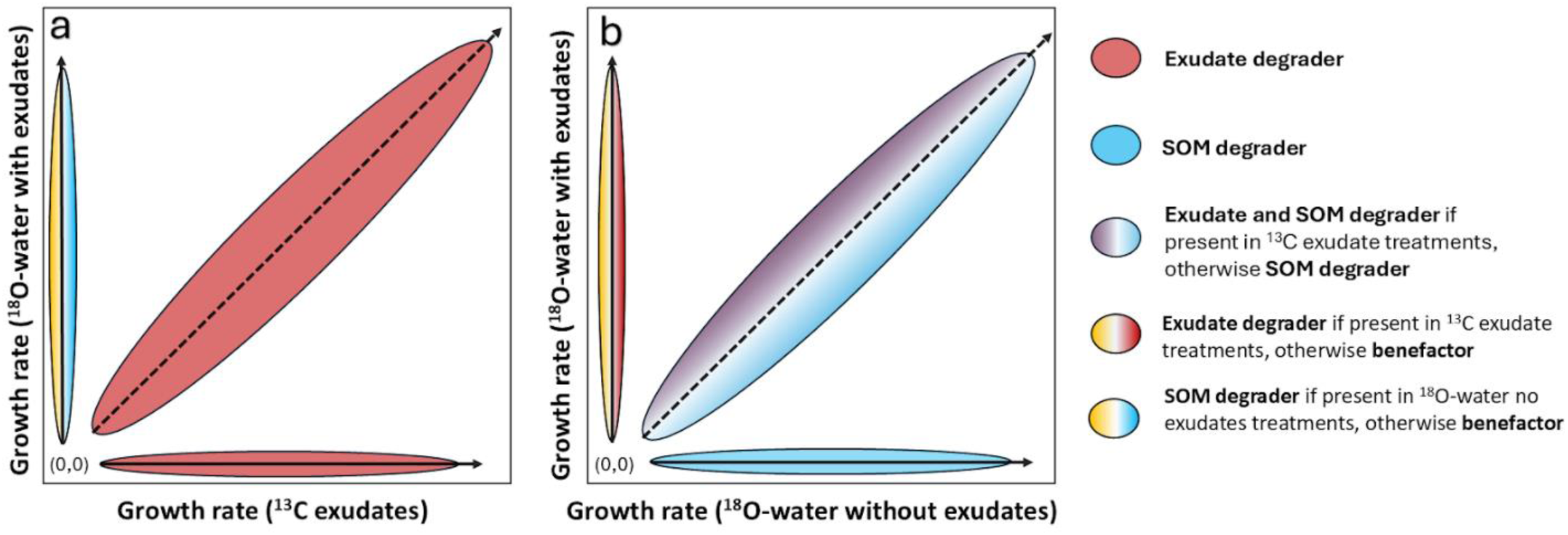
Workflow and strategy overview for interpreting subsequent data. The presence of shared ASVs across exudate addition treatments **(a)** and between the presence and absence of exudate treatments **(b)** reveals distinct microbial strategies represented by colors. Since ^18^O-water with exudates allows for the growth of exudate, SOM and benefactors, ASVs found in this group were determined to be *exudate degraders* if they were also present in ^13^C exudate treatments, and *SOM degraders* if present in in ^18^O-water without exudates treatments. ASV’s present in all three treatments, or between ^18^O-water without exudates and ^13^C exudates were determined to be *exudate and SOM degraders* while *benefactors* were ASVs found only in ^18^O-water with exudates and no other treatments.

We classified microbes into four strategies by plotting their growth rates across treatments: *exudate degraders*, *SOM degraders*, *exudate and SOM degraders*, and *benefactors*. On each plot is a y = x line which represents a theoretical equivalence point where growth rates between the treatments are equal and thus there is no preference for exudates or SOM (Fig. 1). Points that fall on the X or Y axis represent ASVs that were only detected in that treatment and show association towards that treatment’s substrate only. *Exudate degraders* were defined as microbes that exclusively grew under ‘^13^C exudates’ treatments, or grew in both ‘^13^C exudates’ and ‘^18^O-water with exudates’ treatments (Fig. 1, red). *SOM degraders* were classified as microbes that grew exclusively in ‘^18^O-water without exudates’ treatments or grew in ‘^18^O-water with exudates’ but not ‘^13^C exudate’ treatments (Fig. 1, blue). *Exudate and SOM degraders* were microbes that grew in all three treatments, demonstrating uptake of both substrates (Fig 1, purple). *Benefactors* represent microbes that were stimulated by the presence of exudates but did not incorporate exudates into their DNA. These microbes grew in ‘^18^O-water with exudates’ treatments but not in either of the other treatments (Fig. 1, yellow).

### 2.14 Statistical analyses and visualizations

All figures and statistical analyses were conducted in R Statistical Software (v4.2.3; R Core Team 2023). To visualize differences in microbial communities across treatments, we generated ordinations using the *phyloseq* package (v1.42.0; McMurdie and Holmes 2013) in conjunction with *ggplot2* (v.3.4.3; Wickham 2016). To test for statistical differences in microbial community compositions across treatments, we conducted permutational multivariate analysis of variance (PERMANOVA) tests using the adonis2 function in the *vegan* package (v2.6-4; Oksanen et al., 2022). We also tested for the assumption of homogeneity of multivariate dispersions using the betadisper function in *vegan*. All species matrices used Bray-Curtis dissimilarity distances due to the high number of zeros present. The model for PERMANOVA tests was: *distance of bacterial or fungal species matrix ∼ inoculant + exudates, permutations = 999*.

To test for statistical differences in respiration rates and cumulative respiration values across treatments we performed a Welch’s ANOVA test using the stats package in base R (R core team, 2024). This was preferred over standard ANOVA due to unequal variances across treatments. The linear model used was *oneway.test(respiration value ∼ treatment, var.equal = FALSE)* where respiration value was replaced with respiration rates or cumulative respiration and treatment was replaced with inoculant or exudate addition factors. Due to unequal variance across treatments, we used Games Howell tests instead of Tukey for means separation using *rstatix* (v. 0.7.2; Kassambara 2023). Effect sizes were calculated using omega squared statistics due to less biases and better performance calculating with unequal variance across treatments (equation 5, Kroes and Finley 2023).

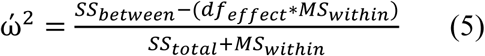

Where SS_between_ represents the sum of squares, or variance between groups, such as the variance between exudate or inoculant treatments, df_effect_ represents the degrees of freedom for the between group being calculated, MS_within_ represents the mean squared variance within the calculated groups, and SS_total_ represents the total variance as the sum of squares (Kassambara 2023).

Heatmaps, to visualize microbial growth rates across treatments, were created using *pheatmap* (v. 1.0.12; Kolde 2019) and *ComplexHeatmap* (v.2.24.1; Gu 2022). Log transformed growth rates (gene copies * gram of dry soil^-1^ * 8 days^-1^) were relativized by the maximum growth rate detected for a given ASV across all treatments.

To test for differences in growth rates across strategies we conducted ANOVA tests followed by Tukey’s Honest Significant Difference means separation tests using the R package *rstatix* (v. 0.7.2; Kassambara 2023). In the case for *Eriophorum* bacterial data, we conducted a Kruskal-Wallis test in base R, followed by a Dunn test in *rstatix* (v. 0.7.2; Kassambara 2023) due to a bimodal distribution of residuals. All residuals were vetted using Q-Q plots and histograms to examine distribution.

## 3. Results

### 3.1 Carbon amendments increase the respiration of rhizosphere inoculated communities but not native permafrost communities

The addition of exudates did not alter cumulative respiration or respiration rates between exudate and no exudate treatments for uninoculated permafrost (Fig. 2a, p-value = 0.771, Table S3, Fig.S1), but did stimulate respiration 3-fold in permafrost inoculated with rhizosphere microorganisms (Fig. 2a, p-value = 6.18e-12, ω² = 0.63, Table S3). Inoculation of permafrost with rhizosphere communities significantly increased cumulative respiration (p-value = 1.27e-10, ω² = 0.45, Table S3) and apparent priming effects (p-value = 4.79e-09, ω² = 0.43, Table S3). There was no difference between *Betula* and *Eriophorum* inoculants for cumulative respiration (Fig. 2a, p-value = 0.978, Table S3), or apparent priming (Fig. 2b, p-value = 0.175, Table S3).

**Fig. 2.**
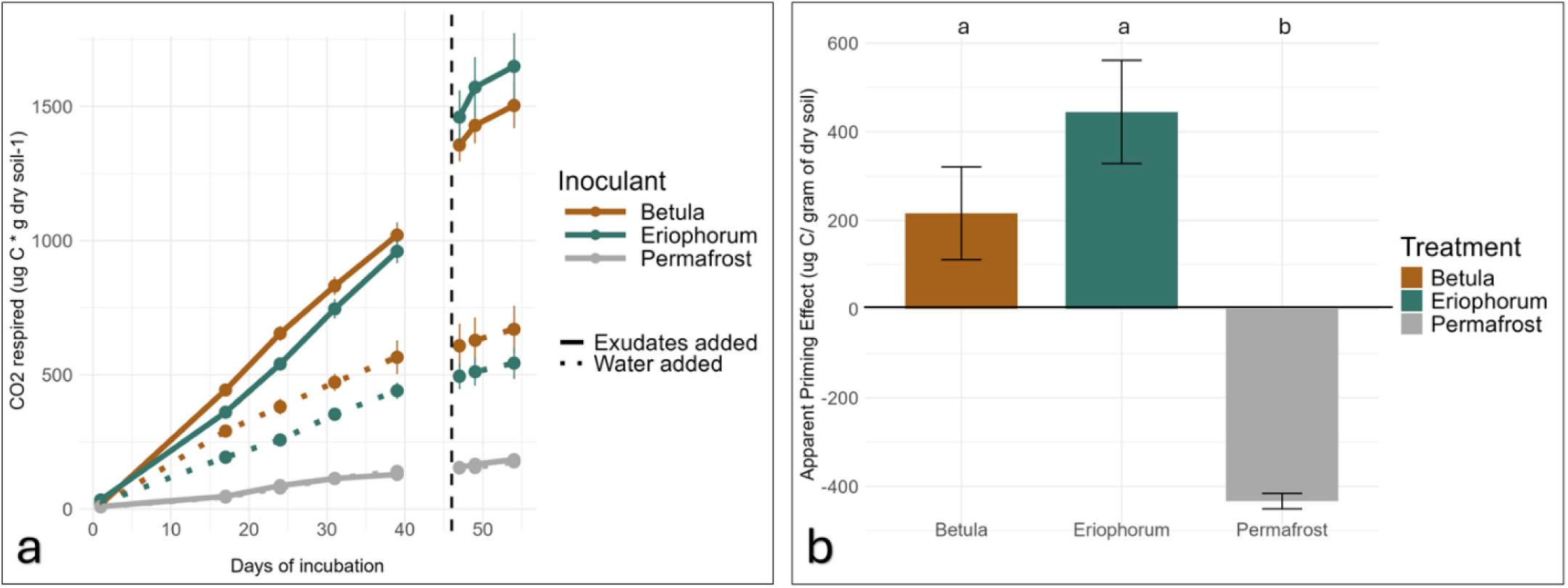
Cumulative respiration **(a)** from an incubation where three inoculants, *Betula* (shrub), *Eriophorum* (sedge), and uninoculated permafrost controls were given simulated root exudates or water daily for 46 days and then spiked with a 7x dose (represented by a vertical dashed line) for the remaining 8 days. Exudate addition resulted in no difference in respiration for uninoculated permafrost treatments (lines are overlapping), whereas *Betula* and *Eriophorum-*inoculated treatments show significantly higher respiration rates when exudates were added. Apparent priming effects throughout the 54 day incubation **(b)** reveal that *Betula* and *Eriophorum*-inoculated permafrost show positive priming while control permafrost communities alone show strong negative apparent priming. Tukey HSD means separation tests reveal inoculant treatments are significantly different from permafrost controls, but not each other, as indicated by their letter designations a-b.

### 3.2 Bacterial and fungal communities show strong inoculant-driven compositional *differences, with bacteria displaying greater overlap across rhizosphere and permafrost environments than fungi*

Inoculating permafrost with rhizosphere communities significantly altered bacterial community structure; both total (including taxa that incorporated the isotope and those that did not; *p* < 0.001, Fig. S2a, Table S3) and active (only taxa that incorporated isotopes; *p* < 0.001, Fig. 3a, Table S3) bacterial communities were significantly affected by inoculation. In contrast, total fungal community structure was unchanged (*p* = 0.289, Fig. S2b, Table S3), whereas active fungal community structure was significantly altered (*p* < 0.001, Fig. 3c, Table S3). Compositionally, 85.3% of active bacterial taxa were found in all inoculants (Fig. 3b), compared to 10.6% of fungal taxa. We detected 468 active bacterial ASV’s across all treatments, 448 of which present in *Betula*, 426 in *Eriophorum*, and 431 in uninoculated permafrost (Fig. 3b). For fungi, we detected a total of 113 unique ASVs across all treatments, 50 of which were detected in *Betula*, 76 in *Eriophorum* and 38 in controls (Fig. 3d). Aside from four taxa, all taxa were shared between exudate and no exudate treatments (Fig. S4).

**Fig. 3.**
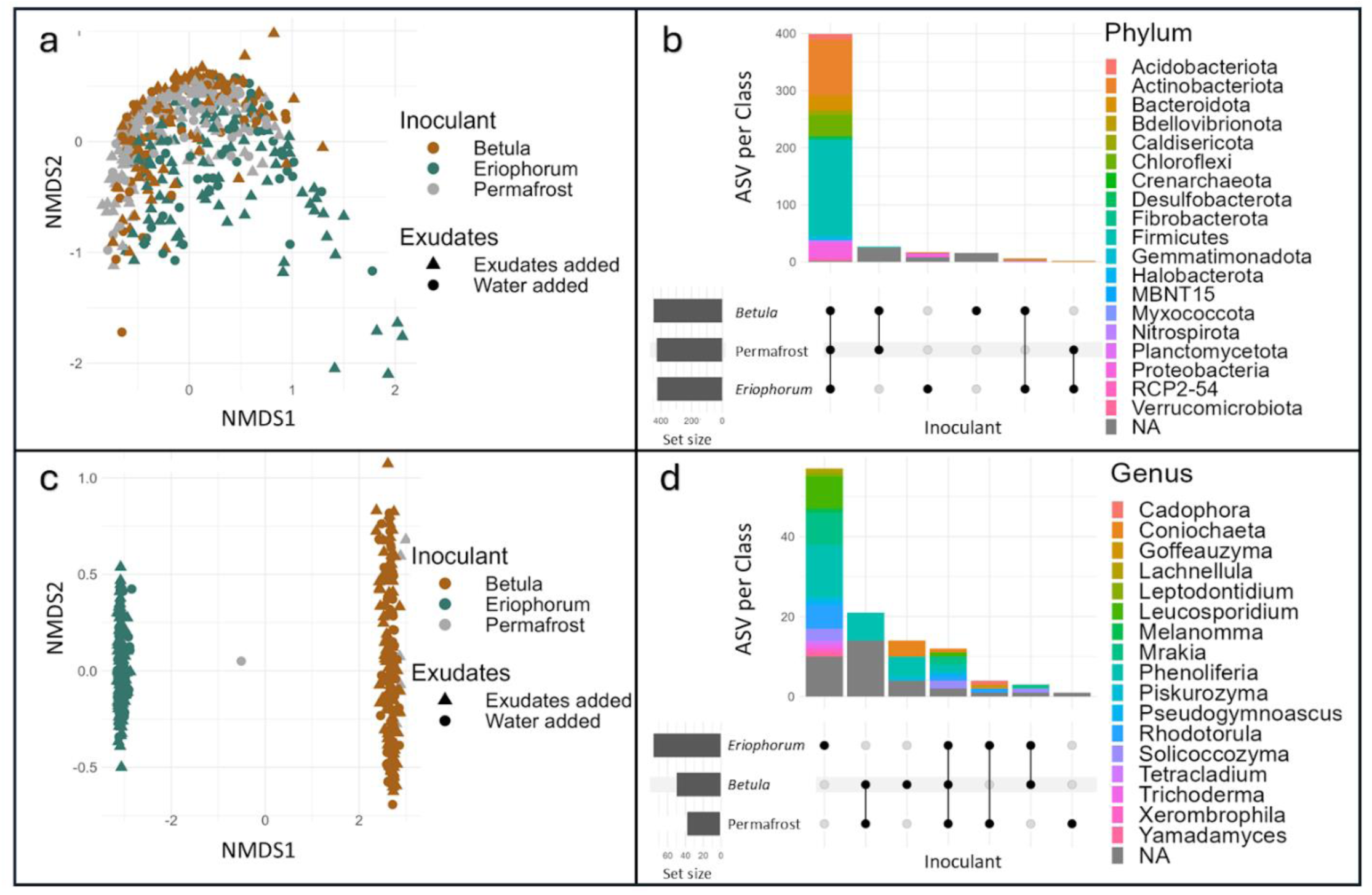
NMDS ordinations of active microbial communities for bacteria **(a)** and fungi **(c)** cluster by inoculants. Upset plots of active communities reveal a high proportion of shared bacterial ASV’s **(b)** present across rhizosphere and permafrost communities with proportionally less shared fungal ASV’s **(d)**. For display purposes we have included only taxa that contained 10 or more reads in our bacterial dataset, and two or more reads in our fungal dataset.

Inoculation altered Shannon diversity due to changes in evenness. Inoculation decreased Shannon diversity for bacteria where uninoculated permafrost diversity was greatest, followed by *Betula* and *Eriophorum-*inoculated communities, respectively (Fig. S5a). Conversely, inoculation increased Shannon diversity in fungal communities and was greatest for *Eriophorum-*inoculated communities followed by *Betula* and uninoculated communities (Fig. S5b). The addition of exudates did not alter Shannon diversity (Fig. S5c, d), nor did it significantly alter bacterial or fungal community structures (p > 0.400, Table S3).

Using qSIP, we quantified taxon-specific growth rates across inoculant communities and exudate conditions and examined patterns in abundance and activity within taxonomic classifications (Fig. 4, S6). Active bacteria were dominated by Actinobacteriota, Bacteroidota, Chloroflexi, and Firmucutes phyla (Fig. 4a, b), and fungal taxa consisted of Phenoliferia genera and various fungi unclassified past kingdom (Fig. 4c, d). Bacterial growth rates with and without exudates were similar for *Betula* and permafrost inoculants, while *Eriophorum*-associated bacteria grew ∼2-10 times faster without exudates (Fig. 4b, S6a, b). Less fungi were detected compared to bacteria with many fungal taxa exhibiting specificity to certain inoculants (Fig. 4c, d, S6c, d).

**Fig. 4.**
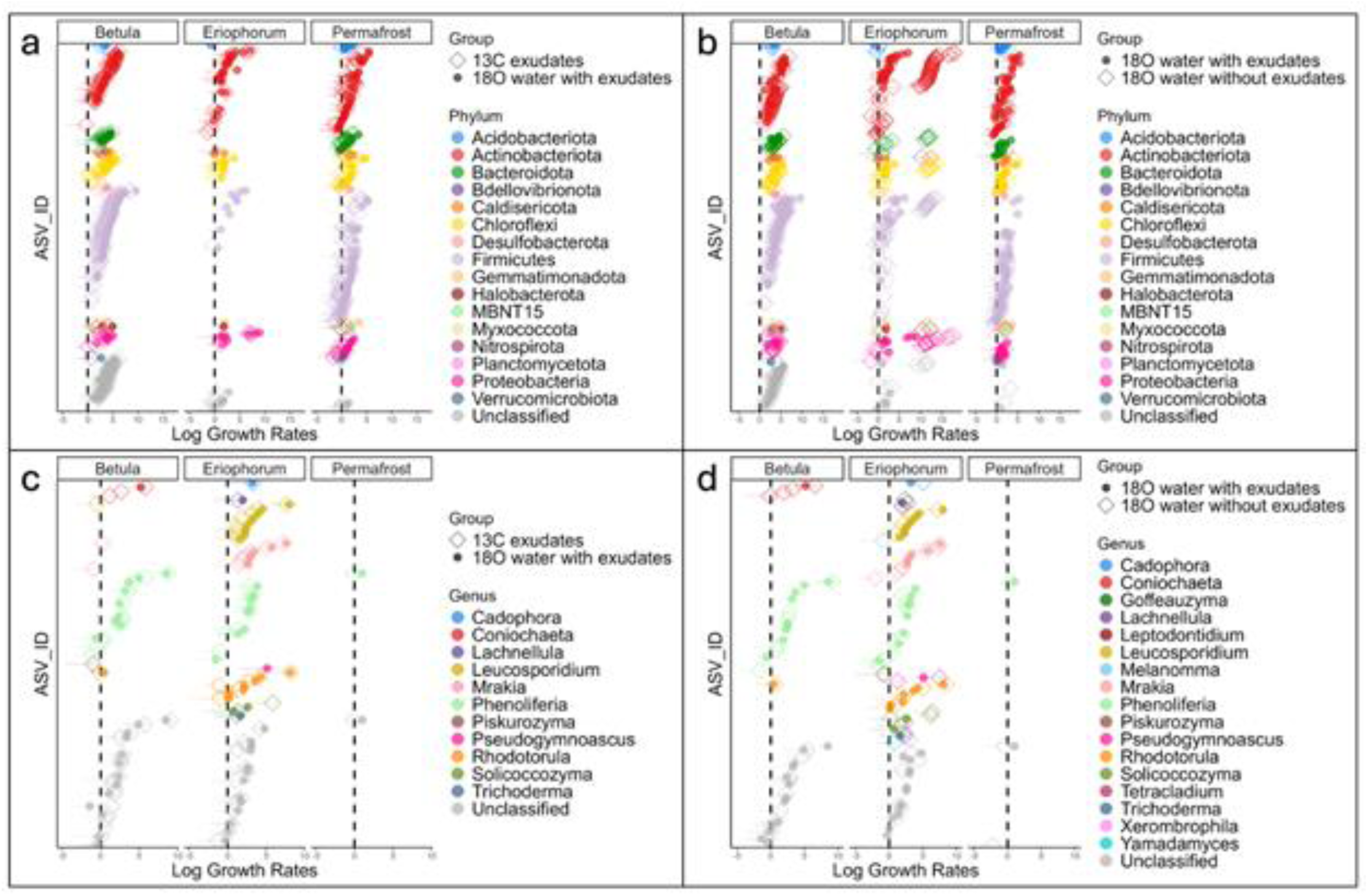
Growth rates calculated from quantitative stable isotope probing reveals active bacteria **(a, b)** and fungi **(c, d)** where bacteria are colored by Phylum and fungi by Genus. Each ASV is fixed along the Y-axis. Growth rates are displayed on the log scale and represent the number of gene copies produced per gram of dry soil over eight days. Columns represent *Betula* and *Eriophorum* rhizosphere communities or uninoculated permafrost communities. Confidence intervals on the tails of points represent the range of calculated growth rates. In the left panels **(a, c)**, exudates are added in both isotope treatments, either as ^13^C exudates or natural abundance exudates with ^18^O-water. The right panels **(b, d)** show the effects of exudate addition, or lack thereof, using growth rates from^18^O-water with exudates and ^18^O-water without exudates.

### 3.3 Patterns in growth rates between treatments indicate differences in substrate preference and strategies

When comparing exudate addition treatments, most *Betula* bacterial taxa were identified as exudate degraders (Fig. 1, Table S2), as they grew in the presence of and consumed exudates, indicated by their positions along the y = x line and X-axis (Fig. 5a). Permafrost bacteria exhibited a similar trend as *Betula* with many classified as *exudate degraders* (Fig. 1, Table S2), as they were also found on the y = x line and X-axis (Fig. 5a). Some permafrost and *Betula* bacteria were classified as *SOM degraders* and *benefactors* (Fig. 1, Table S2), as indicated by their grouping on the Y-axis (Fig. 5a), whereas almost all *Eriophorum* bacteria were *SOM degraders* (Fig. 1, Table S2) as they grouped on Y-axis and were also detected in ^18^O-water without exudates treatments (Fig. 5a).

**Fig. 5.**
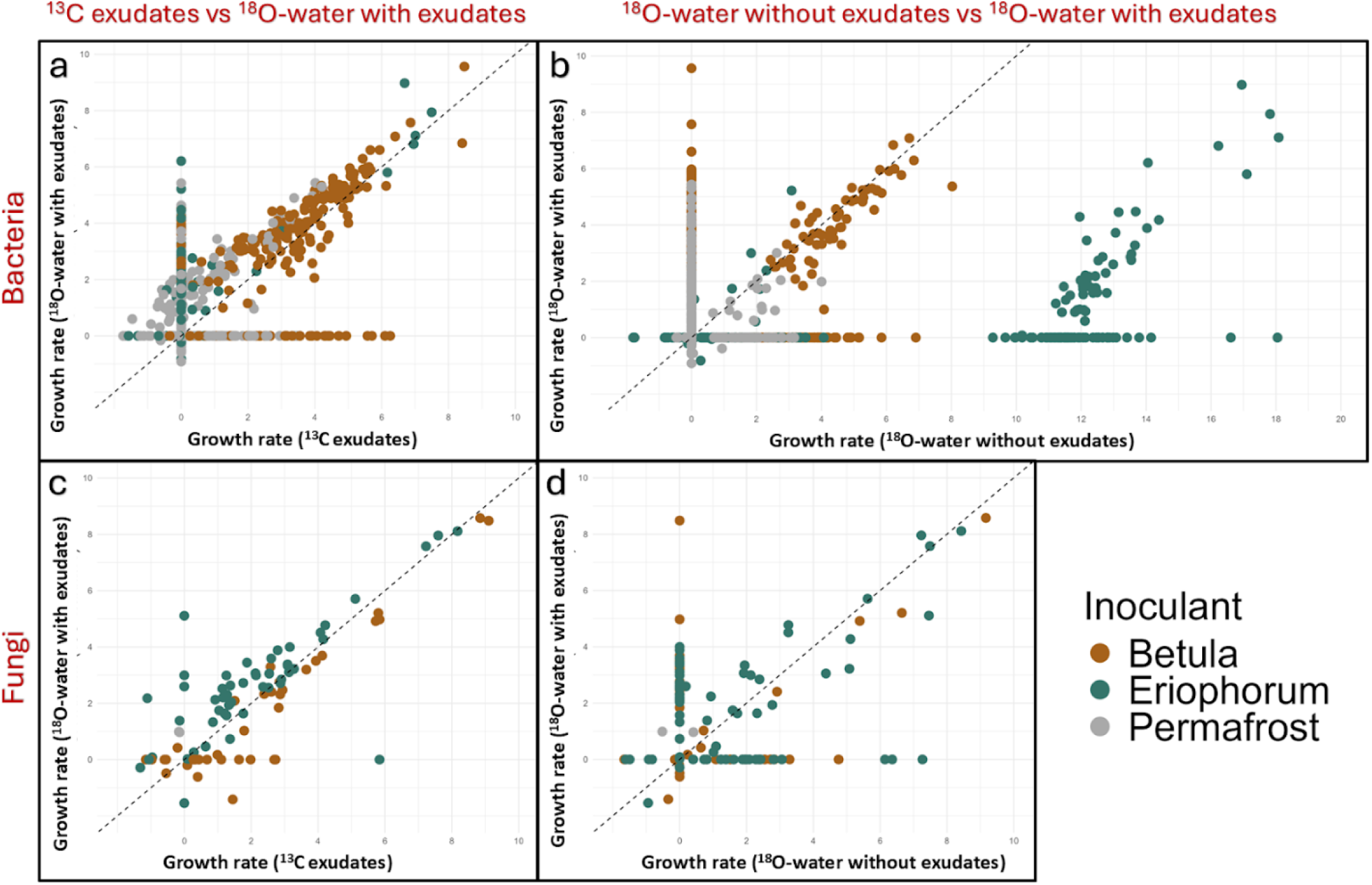
Emergent life history strategies are revealed by combining the log growth rates (gene copies * gram of dry soil^-1^ * 8 days^-1^) of various isotope treatments. Each point represents an ASV where bacterial **(a, b)** and fungal **(c, d)** ASVs are compared across ^13^C and ^18^O exudate treatments **(a, c)** and ^18^O exudate and ^18^O no exudate **(b, d)**. Comparisons between complementary isotope treatments allow for strategy designations (see Fig. 1). Dashed lines at the y = x slope represent the point where the growth rates between the two isotope treatments are equal. Points found on an X or Y axis represent taxa found only in that treatment.

For comparisons between exudate and no exudate bacterial groups, *Betula* were mostly *exudate and SOM degraders* (Fig. 1, Table S2) as they again clustered near the y = x line (Fig. 5b). Permafrost bacteria were primarily classified as *exudate degraders*, *SOM degraders*, or *benefactors* (Fig. 1, Table S2), as they resided mostly on the X and Y axes (Fig. 5b). *Eriophorum* bacteria were largely *exudate and SOM degraders* or *SOM degraders* (Fig. 1, Table S2) as they clustered either on the X-axis or were far right of the y = x line indicating growth in both treatments (Fig. 5b).

Trends in fungal growth rates did not match bacteria. For exudate treatment comparisons, *Betula* and *Eriophorum* fungi were mostly found to be *exudate degraders* (Fig. 1, Table S2) and clustered near the y = x line (Fig. 5c). Some fungi were *SOM* degraders which included all fungi found on the Y-axis (Fig. 5c).

Comparisons between exudate and no exudate groups for fungi revealed a mixture of *exudate degraders*, *exudate and SOM degraders*, and *SOM degraders* (Fig 1, Table S2) as indicated by the spread of taxa across the y = x line and X and Y axes (Fig. 5d). We did not detect any *benefactors* for fungi.

### 3.4 Individual taxa exhibit distinct growth patterns based on community context and exudate additions

We generated heatmaps to examine how growth rates varied for an individual ASV across strategies and inoculants (Fig. 6, S7). We found that growth rates and strategies associated with a given ASV were highly dependent on the inoculant community and exudate conditions (Fig. 6, S7). Similarly, we found that inoculant communities contained unique profiles of bacterial strategists where *Betula* was dominated by *exudate degraders*, *exudate and SOM degraders*, and *SOM degraders*; *Eriophorum* was dominated by *SOM degraders*; and permafrost was dominated by *exudate degraders, SOM degraders,* and *benefactors* (Fig. 6a, Table S2). Profiles for fungal strategists revealed *Betula* was dominated by *exudate degraders* and *exudate and SOM degraders*; whereas *Eriophorum* was dominated by *exudate degraders, exudate and SOM degraders*, and *SOM degraders* (Fig. 6b, Table S2). Uninoculated permafrost had only three detectable fungal taxa, two of which were *exudate and SOM degraders* and one was *SOM degrader* (Fig. 6b, Table S2).

**Fig. 6.**
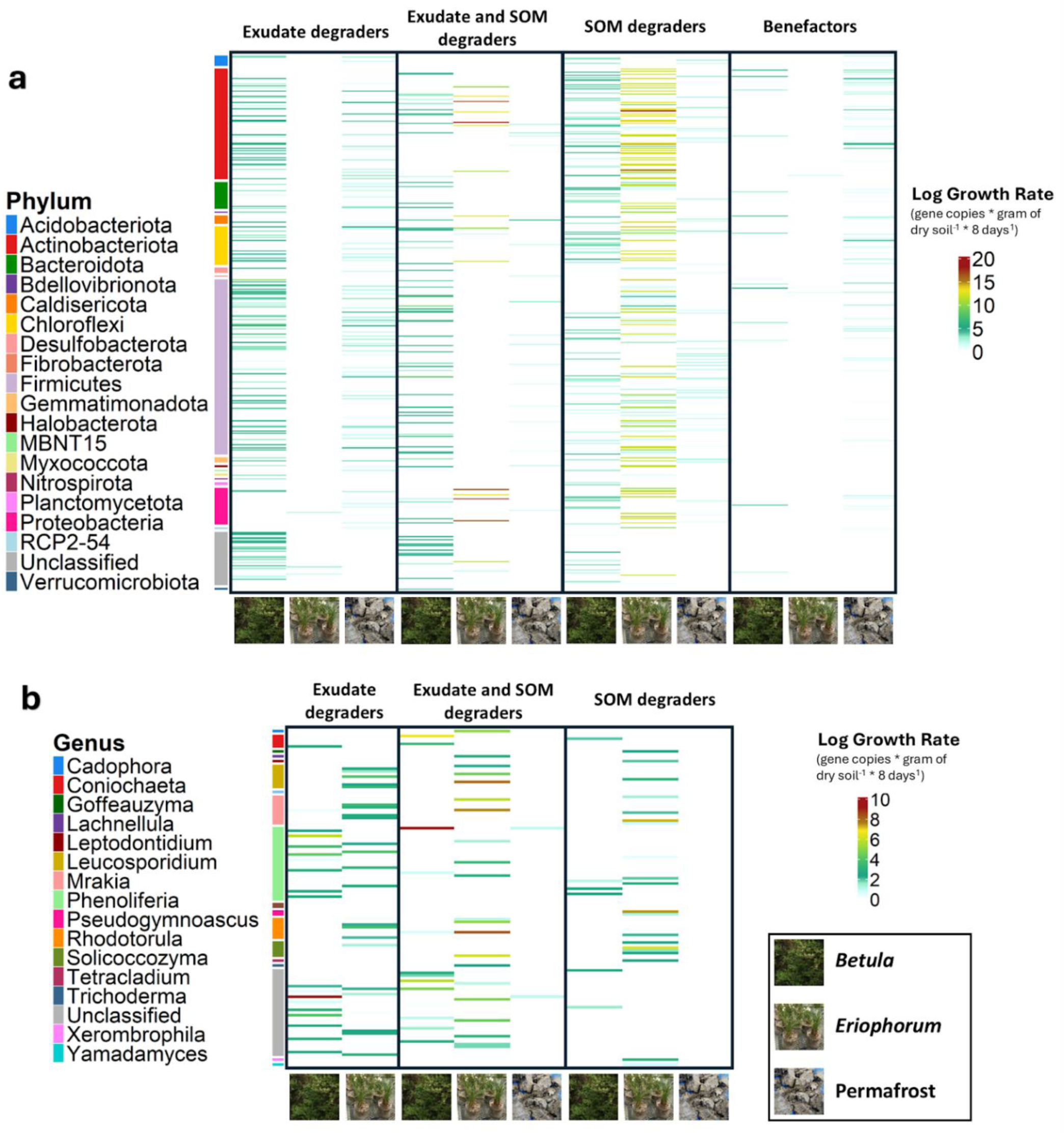
Heat plots reveal distinct patterns of taxon-specific growth rates corresponding to inoculant communities and isotope treatments for bacteria **(a)** and fungi **(b)**. Each row is an individual taxon set in a fixed position across plots. Each column represents an inoculant community (*Betula, Eriophorum* or permafrost). Columns are arranged by microbial strategies (exudate degrader, exudate and SOM degrader, SOM degrader, and benefactor). Growth rates are displayed on a log scale where warmer colors indicate fast growth and cooler indicate slower growth.

### 3.5 Rhizosphere and permafrost microbes display similar median growth rates across strategists at distinct proportions within an inoculant environment

Box plots were generated for bacteria (Fig. 7a) and fungi (Fig. 7c) to examine the distribution of growth rates across strategists and inoculants. Median log transformed growth rates for *Betula* and uninoculated permafrost bacteria were consistent across strategies at ∼3.5 and ∼1.3 gene copies * gram of dry soil^-1^ * 8 days^-1^ respectively, while *Eriophorum* displayed differential growth rates across strategies at ∼ 2.5 gene copies * gram of dry soil^-1^ * 8 days^-1^ for *SOM degraders* and *exudate and SOM degraders* compared to 0.3 for *exudate degraders* and 1.1 for *benefactors* (Fig. 7a). Median growth rates for fungi were consistent across *Betula* and *Eriophorum* environments at ∼2.5 gene copies * gram of dry soil^-1^ * 8 days^-1^ and were barely detectable in uninoculated permafrost (Fig. 7c). We did not detect any fungal *benefactors*, nor did we detect any *exudate degraders* for uninoculated permafrost fungi (Fig. 7c). To examine the relative proportion of strategists within inoculant environments we summed the log growth rates of bacteria (Fig. 7b) and fungi (Fig. 7d). We found that growth rates for *Betula* were dominated by *exudate degraders* and *exudate and SOM degraders* for bacteria and fungi (Fig. 7b, d) while *Eriophorum* was dominated by *SOM degraders* for bacteria (Fig. 7b) and *exudate and SOM degraders* for fungi (Fig. 7d). Uninoculated permafrost was dominated by *exudate degraders* for bacteria (Fig. 7b) and contained barely detectable growth in all strategies for fungi (Fig. 7d).

**Fig. 7.**
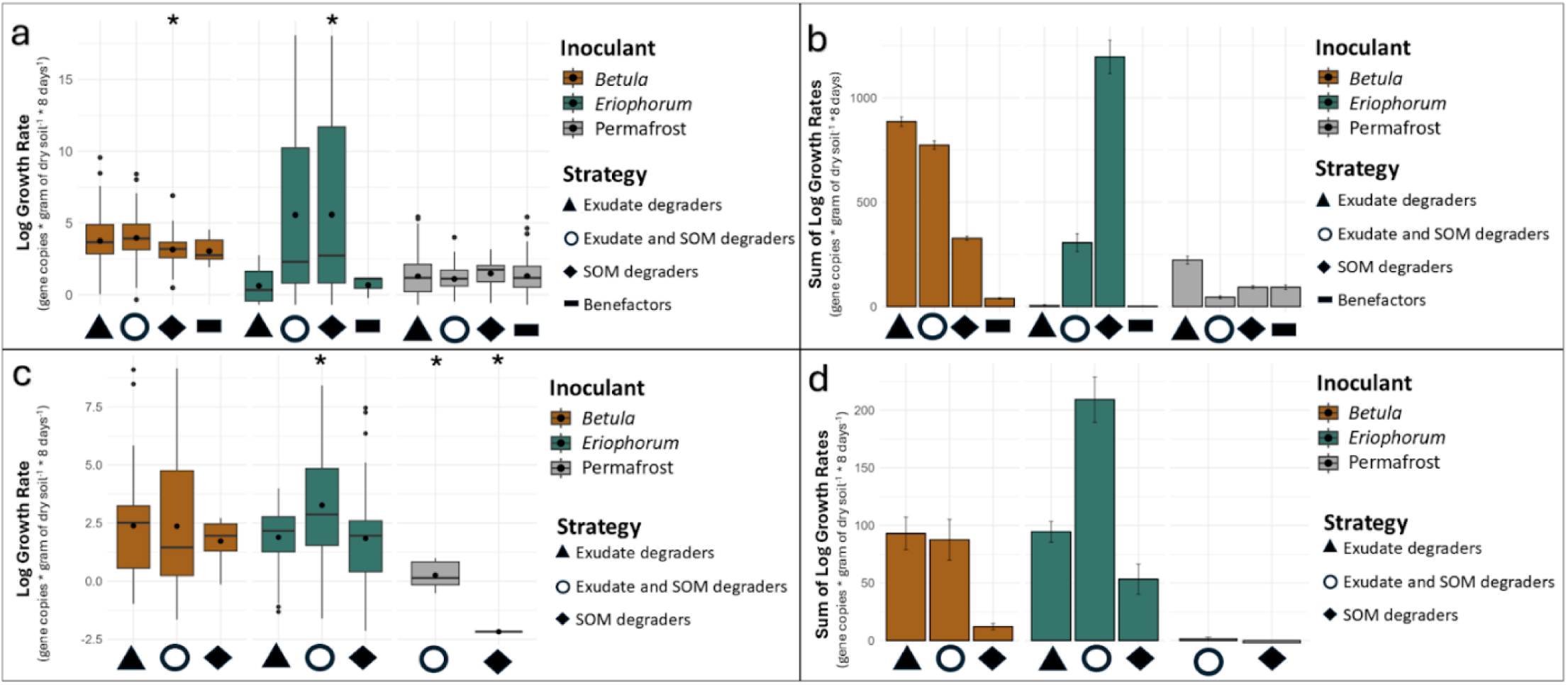
Median growth rates for bacteria **(a)** and fungi **(c)** are shown in box plots on the left, while the sum of growth rates across all ASV’s in an isotope/ exudate treatment are shown in bar graphs on the right for bacteria **(b)** and fungi **(d)**. Strategy refers to metabolic and substrate preferences deduced from comparing presence across multiple isotope and exudate treatments. ‘*’ represents a strategy that has statistically significant growth rates relative to other strategies within an inoculant subset.

## 4. Discussion

The coalescence between rhizosphere and permafrost microbial communities occurs as active layers and rooting depth deepens from permafrost thaw (Blume-Werry et al., 2019, Hewitt et al., 2019, Doherty et al., 2025). Coalescence of these distinct microbial communities are likely to alter tundra carbon cycles by changing the composition and function of the microbial communities in these environments (Parker et al., 2021). Moreover, the presence of root exudates may differentially impact microbial taxa or functional types within rhizosphere communities leading to unique changes in SOM decomposition and metabolic profiles (Blagodatskaya et al., 2021). By performing an incubation experiment inoculating permafrost with rhizosphere communities from the plants *B. nana* and *E. vaginatum*, we were able to examine the influence these distinct rhizosphere communities had on the decomposition of thawed permafrost in the presence and absence of exudates. We found that the addition of exudates onto uninoculated permafrost communities did not result in increased CO_2_ respiration, whereas exudate additions in rhizosphere inoculated communities resulted in significant increases in CO_2_ respiration. Using qSIP with ^13^C and ^18^O isotope solutions embedded in exudate addition treatments, we found all detectable microbial taxa demonstrate a niche based on substrate preference and microbial community context. Finally, we found that *Eriophorum* harbor a disproportionate amount of SOM-targeting bacteria relative to *Betula* and uninoculated permafrost.

### 4.1 The addition of exudates increased soil respiration on rhizosphere-inoculated permafrost but not on uninoculated permafrost

Evidence for permafrost thaw to increase soil respiration has been found in various laboratory incubations (Waldrop et al., 2010, Wild et al., 2014, Ernakovich et al., 2017, Doherty et al., 2020) and field experiments (Trucco et al., 2012, Natali et al., 2015, Blok et al., 2018). The average respiration rate for uninoculated permafrost was ⁓0.2 𝜇g CO_2_-C *grams of dry permafrost^-1^ * hour^-1^ which is in line with other studies that found ⁓0.13-0.20 𝜇g CO_2_-C *gram of dry permafrost^-1^ * hour^-1^ (Waldrop et al., 2010, Wild et al., 2014, Ernakovich et al., 2017, Blok et al., 2018, Doherty et al., 2020). Previous studies have found significant increases in respiration from the amendments of glucose and amino acids (Wild et al., 2014). However, we did not detect a difference in respiration rates in uninoculated permafrost when exudates were added. Because the C:N ratios were comparable between our soils and Wild et al., 2014 at 18 and 21 respectively (7.2% C and 0.4% N (this study); 4.3% C and 0.2% N) it is unlikely that differences in C:N stoichiometry drove respiration patterns.

The low respiration in the presence of exudates in our uninoculated permafrost soils could be due to a variety of factors, including phosphorus limitation, abundance of methanogens or acetogens, the production of toxin or antitoxins, low carbon use efficiency, or low amounts of microbial biomass in permafrost. Phosphorus limitations can stimulate microbes to mobilize phosphorus through dissolution/desorption and hydrolysis of phosphorus sources within SOM, rather than engaging in SOM decomposition (Dijkstra et al., 2013). Alternatively, a high prevalence of methanogens or acetogens could mask the CO_2_ output of the community by incorporating respired CO_2_ into hydrocarbons and microbial products (Kurth et al., 2020). Given the inundation and anoxic areas of both intact and thawed permafrost, the presence of anoxic methanogens and acetogens are common (Mackelprang et al., 2011, Coolen and Orsi 2015, Waldrop et al., 2023). Indeed, we did detect CO_2_ reducing methanogens in our soils, particularly in *Betula* and uninoculated permafrost (Fig. S8). Metagenomic analysis of permafrost from a nearby site, Imnaviat, suggests a high metabolic capability for methanogenesis, acetogenesis, and toxin/ antitoxin production pathways (Waldrop et al., 2023). The production of toxins and antitoxins could also partially explain the lack of increased respiration in response to exudate additions, as toxins and antitoxin production often occurs at the expense of growth and metabolism (Malik et al., 2019). A high carbon use efficiency could also explain low respiration rates in the presence of exudates, as cold adapted microbes tend to have highly efficient enzymes to mitigate energy loss during respiration (Shen et al., 2021). Finally, low amounts of microbial biomass, which are not uncommon in permafrost environments, may also have dampened microbial activity and respiration (Waldrop et al., 2025). Nevertheless, we observed significant increases in respiration in rhizosphere inoculated permafrost and this effect was even more pronounced in rhizosphere-inoculated treatments that received exudates (Fig. 2a), supporting the concept that rhizosphere microbes are a source of CO_2_ respiration and a strong stimulant of SOM decomposition in recently thawed permafrost (Keuper et al., 2020), as well as in active layers of permafrost-affected soils (Street et al., 2020). Our data supports our hypothesis that the intrusion of rhizosphere communities into thawed permafrost increases SOM degradation rates relative to permafrost on its own (Fig. 2b).

### 4.2 Microbial taxa specialize in specific environments based on substrate preference and surrounding microbial communities

Despite the overlap of ASVs in each environment (Fig. 3b, d, S4), differences in relative abundance between inoculants and exudate treatments produced distinct bacterial and fungal communities (Fig. 3a, c, S2, S3). This specificity likely reflects microorganisms’ metabolic specialization, allowing them to thrive in narrow chemical niches while limiting growth in less favorable environments (Jacoby and Kopriva 2019, Schink et al., 2022). We found a handful of taxa that demonstrate this phenomenon. For instance, a bacterium from the genera *Clostridium sensu stricto 13*, was present in both *Betula* and *Eriophorum* rhizosphere inoculants; it grew 10-30 times faster in the *Betula* exudate treatments compared to *Eriophorum* exudate treatments (relative abundance 0.29% and 0.16% respectively) yet grew 2,400 times faster in *Eriophorum* no exudates treatment relative to *Eriophorum* exudate treatments (relative abundance 0.0046% and 0.16% respectively). Likewise, a bacterium from the genera *Robbsia* was present in all *Eriophorum* treatments and grew moderately under exudate additions (relative abundance 0.10%). However, in the absence of exudates the growth rate increased exponentially, between 2,800 and 28,000 times (relative abundance 0.07%). Interestingly, all taxa detected from *Betula* or permafrost environments present in both exudate and no exudate treatments exhibited no preference for one condition over the other, as demonstrated by overlapping confidence intervals on growth rate plots (Fig. 4a, b, S6a, b); whereas *Eriophorum* had many taxa present with and without exudates that grew faster without exudates.

Maximum growth rates for bacteria and fungi are highly dependent on substrates in the environment and microbial community context (Fig. 6, S7). The realized expression of traits represent the intersection between a biochemical niche, abiotic selection pressures, and intra-community interactions (Malard and Guisan 2023). Because abiotic conditions were held constant, our results highlight the importance of intra community dynamics in determining microbial growth rates and substrate preferences. We found that most bacteria and fungi are associated with either exudate or SOM carbon degradation, but their substrate preference can change depending on microbial community context. For example, some bacteria identified as fast-growing *SOM degraders* in *Eriophorum* were moderately-growing *exudate degraders* in *Betula* (Fig. S6), demonstrating metabolic specialization is dependent in part by microbial community dynamics. Despite metabolic flexibility, no ASVs displayed consistently high growth rates across more than one inoculant, suggesting again that substrate preference may be secondary to microbial community context in determining microbial strategies (Fig. 6, S7). Due to the ability for some microbes to switch strategies based on inoculant communities, our second hypothesis, that taxa associated with degrading exudate or SOM carbon would be consistent across treatments, indicating a fixed life history strategy across environments regardless of microbial communities, was not supported.

Our findings are consistent with previous work that suggests emergent growth rates, or growth yield, is dependent on environmental context (Morrison et al., 2022). Although genetic factors, such as codon usage bias and transcriptional efficiency can determine a microbe’s maximum potential growth rate (Weissman et al., 2021, Chuckran et al., 2024), emergent growth rates are dependent on environmental and biochemical conditions which correspond to a microbe’s ability to use available substrates efficiently (Marschmann et al., 2022). One reason why microbes have a maximum growth rate in one community and not the other may be due to the secondary metabolites generated in a particular environment which can result in cross-feeding. For example, respirofermentative bacteria, which prefer sugar substrates, generate organic acids as a secondary metabolite during sugar metabolism. The relative abundance of respirofermentative bacteria influences the downstream production of organic acids and thus the relative abundance of microbes associated with organic acid metabolism (Sun et al., 2024). This example of cross-feeding may explain some of the presence-absence trends in our data. For example, a fungi from the genus *Pheloliferia* was present in all *Betula* and permafrost treatments, yet absent in *Eriophorum*. There was no difference in its growth rate between exudate and no exudate treatments (within the respective *Betula* or uninoculated permafrost treatments) indicating this taxa does not prefer exudates or SOM exclusively. Interestingly, it grew ∼4,500 – 9,000 times faster in *Betula* environments compared to permafrost and did not grow at all in *Eriophorum*. This suggests that the secondary metabolites created by microbes in *Betula* rhizosphere communities drove the heightened proliferation of *Pheloliferia*.

### 4.3 Betula harbors more exudate degraders in the rhizosphere while Eriophorum harbors more SOM degraders

We observed evidence for plant selection of microbial functions (Dennis et al., 2010, Jones et al., 2019). *Betula* and *Eriophorum* possess key distinctions in root architecture, rooting depth, root exudate profiles, association with mycorrhizal fungi, and rhizosphere composition (Wallenstein et al., 2007, Iversen et al., 2015, Schaefer et al., 2025), and these differences are manifestations of their contrasting life history strategies. For example, *Eriophorum* contain rhizomes that grow new vertically extended roots each spring (Wein 1973). Inner roots are less decomposed, allowing dead plant material to account for up to one third of the soil weight beneath the rhizomes (Chapin et al., 1979). Moreover, the ratio of living to dead roots can be as low as 1:45 (Wein 1973), representing a substantial amount of dead plant material surrounding live *Eriophorum* roots (Weintraub and Schimel 2005). The high prevalence of plant necromass could explain why we detected more SOM targeting bacteria in *Eriophorum* communities as opposed to *Betula* and uninoculated permafrost. Through the natural accumulation of SOM around plant roots, *Eriophorum* appears to have selected for *SOM degraders* in its rhizosphere, whereas *Betula,* by forming associations with ectomycorrhizal fungi, has selected against SOM-degrading saprotrophs, catering to a microbial community that is partial to exudate and dissolved organic carbon sources. Thus, our hypothesis that the majority of active taxa would be shared across rhizosphere groups but not permafrost was not supported, as plants display the ability to cater unique compositions of microbial taxa as well as unique proportions of functional types (Fig. 6, 7).

### 4.4 Traditional copiotroph-oligotroph frameworks do not apply to rhizosphere microbes

Most microbial ecological frameworks correlate oligotrophy and SOM degradation with slower growth rates (Chen et al., 2014). Rhizosphere bacteria have been shown to have higher realized growth rates than bacteria in the bulk soil (López et al., 2023). Because the rhizosphere is associated with high rates of carbon exudation, rhizodeposition and nutrient turnover (Jones et al., 2009), they are also described as copiotrophs (López et al., 2023). Our results challenge this traditional framework by finding numerous SOM-degrading taxa in the rhizosphere, particularly *Eriophorum*, that had markedly faster growth rates in environments that were devoid of exudates and rhizodeposits demonstrating that fast growth is not contradictory to an oligotrophic lifestyle (Fig. 4, 5, 7). The observation that SOM-targeting microbial activity is independent of labile carbon inputs is consistent with conceptual frameworks and soil carbon models which articulate nutrient demand is the key driver for SOM degradation in the rhizosphere (Fontaine et al., 2003, Dijkstra et al., 2013, Rocci et al., 2024). Rhizosphere environments support a range of microbial strategies on a continuum between extreme copiotrophy and oligotrophy (Stone et al., 2023). While conceptually copiotrophs are linked to the acquisition of easily metabolized substrates and oligotrophs are linked to the decomposition of a variety of root substrates (Malik et al., 2019; Marschmann et al., 2024), our work shows that for many microbes, their emergent strategy—or realized niche based on substrate preference, substrate availability, and microbe-microbe interactions—is malleable and highly context dependent. We found numerous instances in which an ASV exhibited faster growth rates in exudate treatments than non-exudate treatments under *Betula*, yet produced the opposite effect in *Eriophorum* (Fig. 4, S6). The range in growth rates within a single ASV across environments and within an inoculant community itself highlights the concept that both copiotrophic and oligotrophic lifestyles coexist in the rhizosphere and even within a single organism at different spatial and temporal points. Our results emphasize the importance of integrating microbial strategies and functional types with temporal and environmental contexts, as broad classifications—such as rhizosphere or bulk soil microbes, and oligotrophs or copiotrophs—do not adequately predict an organism’s metabolic output (Morrissey et al., 2019; Malard and Guisan, 2023).

## 5. Conclusion

The tundra is rapidly changing; permafrost thaw and increased shrub growth are altering nutrient and carbon cycles (Sturm et al., 2001, Myers-Smith et al., 2011, Schuur et al., 2022). Contrary to other work, which suggests shrub expansion will induce a loss of SOM (Parker et al., 2015, Parker et al., 2021, Street and Caldararu 2022); our results suggest that an expansion of shrubs may mitigate SOM losses induced by permafrost thaw by concurrently reducing the amount of SOM-degrading bacteria associated with *Eriophorum* rhizosphere communities. We recognize that ecosystem interactions between shrub expansion and soil thermodynamics are complex and recognize the potential for positive feedback loops in which shrub growth increases snowpack and soil warming, causing more decomposition of SOM and shrub growth (Sturm et al., 2005). Likewise, we recognize that long term effects on SOM dynamics, such as the formation of SOM over seasonal and annual timescales, can offset losses in SOM due to decomposition. A meaningful follow up could be to examine SOM formation rates attributed to different rhizosphere communities and microbial taxa. Nevertheless, our work highlights the consideration of microbial traits when predicting the fate of permafrost carbon and suggests that differences in rhizosphere composition between shrubs and sedges, and their affinities for exudates or SOM, represents a critical component for predicting SOM dynamics and CO_2_ fluxes in thawing permafrost systems.

## Acknowledgments

We would like to acknowledge Else Schlerman for help collecting permafrost samples and Nathan Alexander for his help with data collection. We would also like to thank the research staff at Lawrence Livermore National Laboratory, especially Jessica Wollard for her help in collecting and processing qPCR data.

## Data availability

Sequence data is available on NCBI Sequence Read Archive: (BioProject PRJNA1250578). Metadata and code is available on Dryad DOI: 10.5061/dryad.9s4mw6mv8 (pending).

## Declaration of generative AI and AI-assisted technologies in the manuscript preparation process

During the preparation of this work the authors used ChatGTP to assist with code used for data analysis and figure generation. After using this tool, the authors reviewed and edited the content as needed and take full responsibility for the content of the published article.

## Author contributions (CRediT authorship contribution statement)

**Sean R. Schaefer**: Writing – original draft, Conceptualization, Data curation, Formal analysis, Funding acquisition, Investigation, Methodology, Visualization. **Steven J. Blazewicz**: Writing – review and editing, Conceptualization, Funding acquisition, Investigation, Methodology, Project administration, Resources, Supervision, Validation. **Hannah Holland-Moritz**: Writing – review and editing, Conceptualization, Formal analysis, Investigation. **Caitlin Hicks Pries**: Writing – review and editing, Funding acquisition, Investigation. **Mike Allen**: Formal analysis, Methodology, Data curation. **Jennifer Pett-Ridge**: Writing – review and editing, Funding acquisition, Resources, Supervision, Validation. **Fernando Montaño-López**: Writing – review and editing, Investigation, Methodology. **A. Stuart Grandy**: Writing – review and editing, Funding acquisition, Investigation. **William Wieder**: Writing – review and editing, Funding acquisition, Conceptualization. **Jessica Ernakovich:** Writing – review and editing, Funding acquisition, Investigation, Methodology, Project administration, Resources, Supervision, Validation.

## References

Apprill, A., McNally, S.P., Parsons, R.J., Weber, L., 2015. Minor revision to V4 region SSU rRNA 806R gene primer greatly increases detection of SAR11 bacterioplankton. Aquatic Microbial Ecology 75, 129–137.

Black, R.F., 1954. Permafrost: a review. Geological Society of America Bulletin 65, 839–856.

Blagodatskaya, E., Kuzyakov, Y., 2008. Mechanisms of real and apparent priming effects and their dependence on soil microbial biomass and community structure: Critical review. Biology and Fertility of Soils 45, 115–131. doi:10.1007/s00374-008-0334-y

Blagodatskaya, E., Tarkka, M., Knief, C., Koller, R., Peth, S., Schmidt, V., Spielvogel, S., Uteau, D., Weber, M., Razavi, B.S., 2021. Bridging Microbial Functional Traits With Localized Process Rates at Soil Interfaces. Frontiers in Microbiology 12. doi:10.3389/fmicb.2021.625697

Blazewicz, S.J., Hungate, B.A., Koch, B.J., Nuccio, E.E., Morrissey, E., Brodie, E.L., Schwartz, E., Pett-Ridge, J., Firestone, M.K., 2020. Taxon-specific microbial growth and mortality patterns reveal distinct temporal population responses to rewetting in a California grassland soil. The ISME Journal 14, 1520–1532. doi:10.1038/s41396-020-0617-3

Blazewicz, S.J., Schwartz, E., 2011. Dynamics of 18O Incorporation from H218O into Soil Microbial DNA. Microbial Ecology 61, 911–916. doi:10.1007/s00248-011-9826-7

Blok, D., Faucherre, S., Banyasz, I., Rinnan, R., Michelsen, A., Elberling, B., 2018. Contrasting above- and belowground organic matter decomposition and carbon and nitrogen dynamics in response to warming in High Arctic tundra. Global Change Biology 24, 2660–2672. doi:10.1111/gcb.14017

Blume-Werry, G., Milbau, A., Teuber, L.M., Johansson, M., Dorrepaal, E., 2019. Dwelling in the deep – strongly increased root growth and rooting depth enhance plant interactions with thawing permafrost soil. New Phytologist 223, 1328–1339. doi:10.1111/nph.15903

Burkert, A., Douglas, T.A., Waldrop, M.P., Mackelprang, R., 2019. Changes in the Active, Dead, and Dormant Microbial Community Structure across a Pleistocene Permafrost Chronosequence. Applied and Environmental Microbiology 85, e02646–18. doi:10.1128/aem.02646-18

Callahan, B.J., McMurdie, P.J., Rosen, M.J., Han, A.W., Johnson, A.J.A., Holmes, S.P., 2016. DADA2: High-resolution sample inference from Illumina amplicon data. Nature Methods 13, 581–583. doi:10.1038/nmeth.3869

Caporaso, G.J., Ackermann, G., Apprill, A., Bauer, M., Berg-Lyons, D., Betley, J., Fierer, N., Fraser, L., Fuhrman, J.A., Gilbert, J.A., Gormley, N., Humphrey, G., Huntley, J., Jansson, J.K., Knight, R., Lauber, C.L., Lozupone, C.A., McNally, S., Needham, D.M., Owens, S.M., Parada, A.E., Parsons, R., Smith, G.T., Thompson, L., Turnbaugh, P.J., Walters, W.A., Weber, L., 2018. EMP 16S Illumina Amplicon Protocol. Protocols.Io.

Chapin, F.S., van Cleve, K., Chapin, M.C., 1979. Soil Temperature and Nutrient Cycling in the Tussock Growth Form of Eriophorum Vaginatum. Journal of Ecology 67, 169–189. doi:10.2307/2259343

Chen, R., Senbayram, M., Blagodatsky, S., Myachina, O., Dittert, K., Lin, X., Blagodatskaya, E., Kuzyakov, Y., 2014. Soil C and N availability determine the priming effect: microbial N mining and stoichiometric decomposition theories. Global Change Biology 20, 2356–2367. 10.1111/gcb.12475

Chen, Y., Xi, J., Xiao, M., Wang, S., Chen, W., Liu, F., Shao, Y., Yuan, Z., 2022. Soil fungal communities show more specificity than bacteria for plant species composition in a temperate forest in China. BMC Microbiology 22, 208. doi:10.1186/s12866-022-02591-1

Chuckran, P.F., Estera-Molina, K., Nicolas, A.M., Sieradzki, E.T., Dijkstra, P., Firestone, M.K., Pett-Ridge, J., Blazewicz, S.J., 2024. Codon bias, nucleotide selection, and genome size predict in situ bacterial growth rate and transcription in rewetted soil. bioRxiv 2024.06.28.601247. doi:10.1101/2024.06.28.601247

Coolen, M.J.L., Orsi, W.D., 2015. The transcriptional response of microbial communities in thawing Alaskan permafrost soils. Frontiers in Microbiology 6. doi:10.3389/fmicb.2015.00197

Dennis, P.G., Miller, A.J., Hirsch, P.R., 2010. Are root exudates more important than other sources of rhizodeposits in structuring rhizosphere bacterial communities? FEMS Microbiology Ecology 72, 313–327. doi:10.1111/j.1574-6941.2010.00860.x

Dijkstra, F., Carrillo, Y., Pendall, E., Morgan, J., 2013. Rhizosphere priming: a nutrient perspective. Frontiers in Microbiology 4. doi:10.3389/fmicb.2013.00216

Doherty, S.J., Barbato, R.A., Grandy, A.S., Thomas, W.K., Monteux, S., Dorrepaal, E., Johansson, M., Ernakovich, J.G., 2020. The Transition From Stochastic to Deterministic Bacterial Community Assembly During Permafrost Thaw Succession. Frontiers in Microbiology 11. doi:10.3389/fmicb.2020.596589

Doherty, S.J., Thurston, A.K., Barbato, R.A., 2025. Active layer and permafrost microbial community coalescence increases soil activity and diversity in mixed communities compared to permafrost alone. Frontiers in Microbiology Volume 16–2025.

Dunford, E.A., Neufeld, J.D., 2010. DNA stable-isotope probing (DNA-SIP). J Vis Exp. doi:10.3791/2027

Ernakovich, J.G., Lynch, L.M., Brewer, P.E., Calderon, F.J., Wallenstein, M.D., 2017. Redox and temperature-sensitive changes in microbial communities and soil chemistry dictate greenhouse gas loss from thawed permafrost. Biogeochemistry 134, 183–200. doi:10.1007/s10533-017-0354-5

Ernakovich, J.G., Wallenstein, M.D., 2015. Permafrost microbial community traits and functional diversity indicate low activity at in situ thaw temperatures. Soil Biology and Biochemistry 87, 78–89. 10.1016/j.soilbio.2015.04.009

Feng, J., Wang, C., Lei, J., Yang, Y., Yan, Q., Zhou, X., Tao, X., Ning, D., Yuan, M.M., Qin, Y., Shi, Z.J., Guo, X., He, Z., Van Nostrand, J.D., Wu, L., Bracho-Garillo, R.G., Penton, C.R., Cole, J.R., Konstantinidis, K.T., Luo, Y., Schuur, E.A.G., Tiedje, J.M., Zhou, J., 2020. Warming-induced permafrost thaw exacerbates tundra soil carbon decomposition mediated by microbial community. Microbiome 8, 3. doi:10.1186/s40168-019-0778-3

Finley, B., Dijkstra, P., Rasmussen, C., Schwartz, E., Mau, R., Liu, X.-J.A., Van Gestel, N., Hungate, B., 2018. Soil mineral assemblage and substrate quality effects on microbial priming. Geoderma 322, 38–47. doi:10.1016/j.geoderma.2018.01.039

Fontaine, S., Mariotti, A., Abbadie, L., 2003. The priming effect of organic matter: a question of microbial competition? Soil Biology and Biochemistry 35, 837–843. 10.1016/S0038-0717(03)00123-8

Geyer, K.M., Dijkstra, P., Sinsabaugh, R., Frey, S.D., 2019. Clarifying the interpretation of carbon use efficiency in soil through methods comparison. Soil Biology and Biochemistry 128, 79–88. 10.1016/j.soilbio.2018.09.036

Gu, Z., 2022. Complex Heatmap Visualization. iMeta. doi:10.1002/imt2.43

Hewitt, R., DeVan, R., Lagutina, I., Genet, H., McGuire, A., Taylor, D.L., Mack, M., 2019. Mycobiont contribution to tundra plant acquisition of permafrost-derived nitrogen. New Phytologist 226. doi:10.1111/nph.16235

Hicks, L.C., Leizeaga, A., Rousk, K., Michelsen, A., Rousk, J., 2020. Simulated rhizosphere deposits induce microbial N-mining that may accelerate shrubification in the subarctic. Ecology 101, e03094. 10.1002/ecy.3094

Hungate, B.A., Mau, R.L., Schwartz, E., Caporaso, J.G., Dijkstra, P., van Gestel, N., Koch, B.J., Liu, C.M., McHugh, T.A., Marks, J.C., Morrissey, E.M., Price, L.B., 2015. Quantitative Microbial Ecology through Stable Isotope Probing. Applied and Environmental Microbiology 81, 7570–7581. doi:10.1128/aem.02280-15

Iversen, C.M., Sloan, V.L., Sullivan, P.F., Euskirchen, E.S., McGuire, A.D., Norby, R.J., Walker, A.P., Warren, J.M., Wullschleger, S.D., 2015. The unseen iceberg: plant roots in arctic tundra. New Phytologist 205, 34–58. 10.1111/nph.13003

Jacoby, R.P., Kopriva, S., 2019. Metabolic niches in the rhizosphere microbiome: new tools and approaches to analyse metabolic mechanisms of plant–microbe nutrient exchange. Journal of Experimental Botany 70, 1087–1094. doi:10.1093/jxb/ery438

Jones, D.L., Nguyen, C., Finlay, R.D., 2009. Carbon flow in the rhizosphere: carbon trading at the soil–root interface. Plant and Soil 321, 5–33. doi:10.1007/s11104-009-9925-0

Jones, P., Garcia, B.J., Furches, A., Tuskan, G.A., Jacobson, D., 2019. Plant Host-Associated Mechanisms for Microbial Selection. Front Plant Sci 10, 862–862. doi:10.3389/fpls.2019.00862

Kassambara, A., 2023. rstatix: Pipe-Friendly Framework for Basic Statistical Tests. doi:10.32614/CRAN.package.rstatix

Keuper, F., Wild, B., Kummu, M., Beer, C., Blume-Werry, G., Fontaine, S., Gavazov, K., Gentsch, N., Guggenberger, G., Hugelius, G., Jalava, M., Koven, C., Krab, E.J., Kuhry, P., Monteux, S., Richter, A., Shahzad, T., Weedon, J.T., Dorrepaal, E., 2020. Carbon loss from northern circumpolar permafrost soils amplified by rhizosphere priming. Nature Geoscience 13, 560–565. doi:10.1038/s41561-020-0607-0

Kimbrel, J., 2024. qSIP2: qSIP Analysis.

Klindworth, A., Pruesse, E., Schweer, T., Peplies, J., Quast, C., Horn, M., Glöckner, F.O., 2013. Evaluation of general 16S ribosomal RNA gene PCR primers for classical and next-generation sequencing-based diversity studies. Nucleic Acids Research 41, e1–e1. doi:10.1093/nar/gks808

Koch, B.J., McHugh, T.A., Hayer, M., Schwartz, E., Blazewicz, S.J., Dijkstra, P., van Gestel, N., Marks, J.C., Mau, R.L., Morrissey, E.M., Pett-Ridge, J., Hungate, B.A., 2018. Estimating taxon-specific population dynamics in diverse microbial communities. Ecosphere 9, e02090. doi:10.1002/ecs2.2090

Kolde, R., 2019. Pheatmap.

Kroes, A.D.A., Finley, J.R., 2023. Demystifying omega squared: Practical guidance for effect size in common analysis of variance designs. Psychological Methods. doi:10.1037/met0000581

Kurth, J.M., Op den Camp, H.J.M., Welte, C.U., 2020. Several ways one goal—methanogenesis from unconventional substrates. Applied Microbiology and Biotechnology 104, 6839–6854. doi:10.1007/s00253-020-10724-7

Kuzyakov, Y., 2006. Sources of CO2 efflux from soil and review of partitioning methods. Soil Biology and Biochemistry 38, 425–448. doi:10.1016/j.soilbio.2005.08.020

López, J.L., Fourie, A., Poppeliers, S.W.M., Pappas, N., Sánchez-Gil, J.J., de Jonge, R., Dutilh, B.E., 2023. Growth rate is a dominant factor predicting the rhizosphere effect. The ISME Journal 17, 1396–1405. doi:10.1038/s41396-023-01453-6

Lynch, L.M., Machmuller, M.B., Cotrufo, M.F., Paul, E.A., Wallenstein, M.D., 2018. Tracking the fate of fresh carbon in the Arctic tundra: Will shrub expansion alter responses of soil organic matter to warming? Soil Biology and Biochemistry 120, 134–144. 10.1016/j.soilbio.2018.02.002

Mackelprang, R., Burkert, A., Haw, M., Mahendrarajah, T., Conaway, C.H., Douglas, T.A., Waldrop, M.P., 2017. Microbial survival strategies in ancient permafrost: insights from metagenomics. The ISME Journal 11, 2305–2318. doi:10.1038/ismej.2017.93

Mackelprang, R., Waldrop, M.P., DeAngelis, K.M., David, M.M., Chavarria, K.L., Blazewicz, S.J., Rubin, E.M., Jansson, J.K., 2011. Metagenomic analysis of a permafrost microbial community reveals a rapid response to thaw. Nature 480, 368–371. doi:10.1038/nature10576

Malard, L.A., Guisan, A., 2023. Into the microbial niche. Trends in Ecology & Evolution 38, 936–945. doi:10.1016/j.tree.2023.04.015

Malik, A.A., Martiny, J.B.H., Brodie, E.L., Martiny, A.C., Treseder, K.K., Allison, S.D., 2019. Defining trait-based microbial strategies with consequences for soil carbon cycling under climate change. The ISME Journal. doi:10.1038/s41396-019-0510-0

Marschmann, G.L., Tang, J., Zhalnina, K., Karaoz, U., Cho, H., Le, B., Pett-Ridge, J., Brodie, E.L., 2024. Predictions of rhizosphere microbiome dynamics with a genome-informed and trait-based energy budget model. Nature Microbiology 9, 421–433. doi:10.1038/s41564-023-01582-w

Marschmann, G.L., Tang, J., Zhalnina, K., Karaoz, U., Cho, H., Le, B., Pett-Ridge, J., Brodie, E.L., 2022. Life history strategies and niches of soil bacteria emerge from interacting thermodynamic, biophysical, and metabolic traits. bioRxiv 2022.06.29.498137. doi:10.1101/2022.06.29.498137

Mauritz, M., Bracho, R., Celis, G., Hutchings, J., Natali, S.M., Pegoraro, E., Salmon, V.G., Schädel, C., Webb, E.E., Schuur, E.A.G., 2017. Nonlinear CO2 flux response to 7 years of experimentally induced permafrost thaw. Global Change Biology 23, 3646–3666. doi:10.1111/gcb.13661

McDonald, M.D., Owusu-Ansah, C., Ellenbogen, J.B., Malone, Z.D., Ricketts, M.P., Frolking, S.E., Ernakovich, J.G., Ibba, M., Bagby, S.C., Weissman, J.L., 2024. What is microbial dormancy? Trends in Microbiology 32, 142–150. doi:10.1016/j.tim.2023.08.006

McMurdie, P.J., Holmes, S., 2013. phyloseq: An R package for reproducible interactive analysis and graphics of microbiome census data. PLoS ONE 8, e61217.

Mekonnen, Z.A., Riley, W.J., Berner, L.T., Bouskill, N.J., Torn, M.S., Iwahana, G., Breen, A.L., Myers-Smith, I.H., Criado, M.G., Liu, Y., Euskirchen, E.S., Goetz, S.J., Mack, M.C., Grant, R.F., 2021. Arctic tundra shrubification: a review of mechanisms and impacts on ecosystem carbon balance. Environmental Research Letters 16, 053001. doi:10.1088/1748-9326/abf28b

Morrison, E.W., Whitney, S.A., Geyer, K.M., Sevigny, J.L., Grandy, A.S., Thomas, W.K., DeAngelis, K.M., Frey, S.D., 2022. Evidence for a genetic basis in functional trait tradeoffs with microbial growth rate but not growth yield. Soil Biology and Biochemistry 172, 108765. doi:10.1016/j.soilbio.2022.108765

Morrissey, E.M., Mau, R.L., Hayer, M., Liu, X.-J.A., Schwartz, E., Dijkstra, P., Koch, B.J., Allen, K., Blazewicz, S.J., Hofmockel, K., Pett-Ridge, J., Hungate, B.A., 2019. Evolutionary history constrains microbial traits across environmental variation. Nature Ecology & Evolution 3, 1064–1069. doi:10.1038/s41559-019-0918-y

Myers-Smith, I.H., Forbes, B.C., Wilmking, M., Hallinger, M., Lantz, T., Blok, D., Tape, K.D., Macias-Fauria, M., Sass-Klaassen, U., Lévesque, E., Boudreau, S., Ropars, P., Hermanutz, L., Trant, A., Collier, L.S., Weijers, S., Rozema, J., Rayback, S.A., Schmidt, N.M., Schaepman-Strub, G., Wipf, S., Rixen, C., Ménard, C.B., Venn, S., Goetz, S., Andreu-Hayles, L., Elmendorf, S., Ravolainen, V., Welker, J., Grogan, P., Epstein, H.E., Hik, D.S., 2011. Shrub expansion in tundra ecosystems: dynamics, impacts and research priorities. Environmental Research Letters 6, 045509. doi:10.1088/1748-9326/6/4/045509

Natali, S.M., Schuur, E.A.G., Mauritz, M., Schade, J.D., Celis, G., Crummer, K.G., Johnston, C., Krapek, J., Pegoraro, E., Salmon, V.G., Webb, E.E., 2015. Permafrost thaw and soil moisture driving CO2 and CH4 release from upland tundra. Journal of Geophysical Research: Biogeosciences 120, 525–537. 10.1002/2014JG002872

Nuccio, E.E., Blazewicz, S.J., Lafler, M., Campbell, A.N., Kakouridis, A., Kimbrel, J.A., Wollard, J., Vyshenska, D., Riley, R., Tomatsu, A., Hestrin, R., Malmstrom, R.R., Firestone, M., Pett-Ridge, J., 2022. HT-SIP: a semi-automated stable isotope probing pipeline identifies cross-kingdom interactions in the hyphosphere of arbuscular mycorrhizal fungi. Microbiome 10, 199. doi:10.1186/s40168-022-01391-z

Oksanen, J., Simpson, G.L., Blanchet, F.G., Kindt, R., Legendre, P., Minchin, P.R., O’Hara, R.B., Solymos, P., Stevens, M.H.H., Szoecs, E., Wagner, H., Barbour, M., Bedward, M., Bolker, B., Borcard, D., Carvalho, G., Chirico, M., Caceres, M.D., Durand, S., Evangelista, H.B.A., FitzJohn, R., Friendly, M., Furneaux, B., Hannigan, G., Hill, M.O., Lahti, L., McGlinn, D., Ouellette, M.-H., Cunha, E.R., Smith, T., Stier, A., Braak, C.J.F.T., Weedon, J., 2022. vegan: Community Ecology Package.

Parada, A.E., Needham, D.M., Fuhrman, J.A., 2016. Every base matters: assessing small subunit rRNA primers for marine microbiomes with mock communities, time series and global field samples. Environmental Microbiology 18, 1403–1414. doi:10.1111/1462-2920.13023

Parker, T.C., Subke, J.-A., Wookey, P.A., 2015. Rapid carbon turnover beneath shrub and tree vegetation is associated with low soil carbon stocks at a subarctic treeline. Global Change Biology 21, 2070–2081. doi:10.1111/gcb.12793

Parker, T.C., Thurston, A.M., Raundrup, K., Subke, J.-A., Wookey, P.A., Hartley, I.P., 2021. Shrub expansion in the Arctic may induce large-scale carbon losses due to changes in plant-soil interactions. Plant and Soil. doi:10.1007/s11104-021-04919-8

Pörtner, H.-O., Roberts, D.C., Masson-Delmotte, V., Zhai, P., Tignor, M., Poloczanska, E., Mintenbeck, K., Alegría, A., Nicolai, M., Okem, A., Petzold, J., Rama, B., Weyer, N.M., 2019. Summary for Policymakers. In: IPCC Special Report on the Ocean and Cryosphere in a Changing Climate.

R Core Team, 2024. R: A Language and Environment for Statistical Computing. R Foundation for Statistical Computing, Vienna, Austria.

Rantanen, M., Karpechko, A.Y., Lipponen, A., Nordling, K., Hyvärinen, O., Ruosteenoja, K., Vihma, T., Laaksonen, A., 2022. The Arctic has warmed nearly four times faster than the globe since 1979. Communications Earth & Environment 3, 168. doi:10.1038/s43247-022-00498-3

Rillig, M.C., Mansour, I., 2017. Microbial Ecology: Community Coalescence Stirs Things Up. Current Biology 27, R1280–R1282. doi:10.1016/j.cub.2017.10.027

Rivers, A., Weber, K., Gardner, T., Liu, S., Armstrong, S., 2018. ITSxpress: Software to rapidly trim internally transcribed spacer sequences with quality scores for marker gene analysis [version 1; peer review: 2 approved]. F1000Research 7. doi:10.12688/f1000research.15704.1

Rocci, K.S., Cleveland, C.C., Eastman, B.A., Georgiou, K., Grandy, A.S., Hartman, M.D., Hauser, E., Holland-Moritz, H., Kyker-Snowman, E., Pierson, D., Reich, P.B., Schlerman, E.P., Wieder, W.R., 2024. Aligning theoretical and empirical representations of soil carbon-to-nitrogen stoichiometry with process-based terrestrial biogeochemistry models. Soil Biology and Biochemistry 189, 109272. doi:10.1016/j.soilbio.2023.109272

Schaefer, S.R., Montaño-López, F., Holland-Moritz, H., Hicks Pries, C., Ernakovich, J.G., 2025. Rhizosphere Bacteria and Fungi are Differentially Structured by Host Plants, Soil Mineralogy and Ectomycorrhizal Communities in the Alaskan Tundra. Arctic Science. doi:10.1139/as-2025-0012

Scharlemann, J.P.W., Tanner, E.V.J., Hiederer, R., Kapos, V., 2014. Global soil carbon: understanding and managing the largest terrestrial carbon pool. Carbon Management 5, 81–91. doi:10.4155/cmt.13.77

Schink, S.J., Christodoulou, D., Mukherjee, A., Athaide, E., Brunner, V., Fuhrer, T., Bradshaw, G.A., Sauer, U., Basan, M., 2022. Glycolysis/gluconeogenesis specialization in microbes is driven by biochemical constraints of flux sensing. Molecular Systems Biology 18, e10704. doi:10.15252/msb.202110704

Schuur, E.A.G., Abbott, B.W., Commane, R., Ernakovich, J., Euskirchen, E., Hugelius, G., Grosse, G., Jones, M., Koven, C., Leshyk, V., Lawrence, D., Loranty, M.M., Mauritz, M., Olefeldt, D., Natali, S., Rodenhizer, H., Salmon, V., Schädel, C., Strauss, J., Treat, C., Turetsky, M., 2022. Permafrost and Climate Change: Carbon Cycle Feedbacks From the Warming Arctic. Annual Review of Environment and Resources 47, 343–371. doi:10.1146/annurev-environ-012220-011847

Schuur, E.A.G., McGuire, A.D., Schädel, C., Grosse, G., Harden, J.W., Hayes, D.J., Hugelius, G., Koven, C.D., Kuhry, P., Lawrence, D.M., Natali, S.M., Olefeldt, D., Romanovsky, V.E., Schaefer, K., Turetsky, M.R., Treat, C.C., Vonk, J.E., 2015. Climate change and the permafrost carbon feedback. Nature 520, 171–179. doi:10.1038/nature14338

Shen, L., Zhang, S., Chen, G., 2021. Regulated strategies of cold-adapted microorganisms in response to cold: a review. Environmental Science and Pollution Research 28, 68006–68024. doi:10.1007/s11356-021-16843-6

Stone, B.W.G., Dijkstra, P., Finley, B.K., Fitzpatrick, R., Foley, M.M., Hayer, M., Hofmockel, K.S., Koch, B.J., Li, J., Liu, X.J.A., Martinez, A., Mau, R.L., Marks, J., Monsaint-Queeney, V., Morrissey, E.M., Propster, J., Pett-Ridge, J., Purcell, A.M., Schwartz, E., Hungate, B.A., 2023. Life history strategies among soil bacteria—dichotomy for few, continuum for many. The ISME Journal 17, 611–619. doi:10.1038/s41396-022-01354-0

Street, L.E., Caldararu, S., 2022. Why are Arctic shrubs becoming more nitrogen limited? New Phytologist 233, 585–587. doi:10.1111/nph.17841

Street, L.E., Garnett, M.H., Subke, J.-A., Baxter, R., Dean, J.F., Wookey, P.A., 2020. Plant carbon allocation drives turnover of old soil organic matter in permafrost tundra soils. Global Change Biology 26, 4559–4571. 10.1111/gcb.15134

Sturm, M., Douglas, T., Racine, C., Liston, G.E., 2005. Changing snow and shrub conditions affect albedo with global implications. Journal of Geophysical Research: Biogeosciences 110. 10.1029/2005JG000013

Sturm, M., Racine, C., Tape, K., 2001. Increasing shrub abundance in the Arctic. Nature 411, 546–547. doi:10.1038/35079180

Taylor, L.D., Walters, William A., Lennon, Niall J., Bochicchio, James, Krohn, Andrew, Caporaso, J. Gregory, Pennanen, Taina, 2016. Accurate Estimation of Fungal Diversity and Abundance through Improved Lineage-Specific Primers Optimized for Illumina Amplicon Sequencing. Applied and Environmental Microbiology 82, 7217–7226. doi:10.1128/AEM.02576-16

Teixeira, C., Holland-Moritz, H., Granada, C., Bayer, C., Sausen, T., Tonial, F., Petry, C., Frey, S., 2024. Land management of formerly subtropical Atlantic Forest reduces soil carbon stocks and alters microbial community structure and function. Applied Soil Ecology. doi:10.1016/j.apsoil.2023.105252

Trucco, C., Schuur, E.A.G., Natali, S.M., Belshe, E.F., Bracho, R., Vogel, J., 2012. Seven-year trends of CO2exchange in a tundra ecosystem affected by long-term permafrost thaw. Journal of Geophysical Research: Biogeosciences 117. doi:10.1029/2011JG001907

Waldrop, M.P., Chabot, C.L., Liebner, S., Holm, S., Snyder, M.W., Dillon, M., Dudgeon, S.R., Douglas, T.A., Leewis, M.-C., Walter Anthony, K.M., McFarland, J.W., Arp, C.D., Bondurant, A.C., Taş, N., Mackelprang, R., 2023. Permafrost microbial communities and functional genes are structured by latitudinal and soil geochemical gradients. The ISME Journal 17, 1224–1235. doi:10.1038/s41396-023-01429-6

Waldrop, M.P., Ernakovich, J.G., Vishnivetskaya, T.A., Schaefer, S.R., Mackelprang, R., Barta, J., O′Brien, J.M., Winkel, M., Barbato, R.A., Heffernan, L., Leewis, M.-C., Hewitt, R.E., Hultman, J., Sun, Y., Biasi, C., Bradley, J.A., Liebner, S., Ricketts, M.P., Muscarella, M.E., Schütte, U., Abuah, F., Whalen, E., Timling, I., Voigt, C., Taş, N., Lloyd, K.G., Siljanen, H.M.P., Rivkina, E.M., Voříšková, J., Tao, J., Liang, R., Li, Z., Lennon, J.T., Onstott, T.C., 2025. Microbial Ecology of Permafrost Soils: Populations, Processes, and Perspectives. Permafrost and Periglacial Processes n/a. doi:10.1002/ppp.2264

Waldrop, M.P., Wickland, K.P., White, R., Berhe, A.A., Harden, J.W., Romanovsky, V.E., 2010. Molecular investigations into a globally important carbon pool: Permafrost-protected carbon in Alaskan soils. Global Change Biology 16, 2543–2554. doi:10.1111/j.1365-2486.2009.02141.x

Walker, M.D., Walker, D.A., Auerbach, N.A., 1994. Plant communities of a tussock tundra landscape in the Brooks Range Foothills, Alaska. Journal of Vegetation Science 5, 843–866.

Wallenstein, M.D., McMahon, S., Schimel, J., 2007. Bacterial and fungal community structure in Arctic tundra tussock and shrub soils. FEMS Microbiol Ecol 59, 428–35. doi:10.1111/j.1574-6941.2006.00260.x

Wein, R.W., 1973. Eriophorum Vaginatum L. Journal of Ecology 61, 601–615. doi:10.2307/2259047

Weintraub, M.N., Schimel, J.P., 2005. Nitrogen Cycling and the Spread of Shrubs Control Changes in the Carbon Balance of Arctic Tundra Ecosystems. BioScience 55, 408–415. doi:10.1641/0006-3568(2005)055[0408:Ncatso]2.0.Co;2

Weissman, J.L., Hou, S., Fuhrman, J.A., 2021. Estimating maximal microbial growth rates from cultures, metagenomes, and single cells via codon usage patterns. Proceedings of the National Academy of Sciences 118, e2016810118. doi:10.1073/pnas.2016810118

Wickham, H., 2016. ggplot2: Elegant Graphics for Data Analysis. Springer-Verlag New York.

Wild, B., Schnecker, J., Alves, R.J.E., Barsukov, P., Bárta, J., Čapek, P., Gentsch, N., Gittel, A., Guggenberger, G., Lashchinskiy, N., Mikutta, R., Rusalimova, O., Šantrůčková, H., Shibistova, O., Urich, T., Watzka, M., Zrazhevskaya, G., Richter, A., 2014. Input of easily available organic C and N stimulates microbial decomposition of soil organic matter in arctic permafrost soil. Soil Biology and Biochemistry 75, 143–151. 10.1016/j.soilbio.2014.04.014

