## Supplemental materials for "Shrub and Sedge Rhizosphere Communities Display Distinct Affinities Toward Exudates and Soil Organic Matter Degradation: a Quantitative Stable Isotope Probing Analysis"

**Supplemental Information**

**Table S1.** Experimental design table embedded within inoculant treatments

| Description | Duration ID | Duration (days) | Substrate added | Isotope added | # of replicates |
|---|---|---|---|---|---|
| Inoculant controls | $T_0$ | 0 | Water | n/a | 3 |
| Pre-isotope exudates | $T_1$ | 46 | Exudates | n/a | 3 |
| Pre-isotope control | $T_1$ | 46 | Water | n/a | 3 |
| Isotope treatment 1 | $T_{full}$ | 54 | Exudates | n/a | 4 |
| Isotope treatment 2 | $T_{full}$ | 54 | Water | n/a | 4 |
| Isotope treatment 3 | $T_{full}$ | 54 | Exudates | $^{18}O$-water | 4 |
| Isotope treatment 4 | $T_{full}$ | 54 | Water | $^{18}O$-water | 4 |
| Isotope treatment 5 | $T_{full}$ | 54 | Exudates | $^{13}C$ exudates | 4 |

Embedded within each inoculant treatment (*Betula, Eriophorum* and permafrost) were different isotope and exudate conditions, as well as different timepoints were T0 represents uninoculated soils at the start of the experiment and T1 represents soils that were pulled prior to isotope additions. Exudates added refers to whether samples were given synthetic exudate solutions daily where Y is yes and N is no. Isotope added refers to isotopically enriched sources or n/a if there was no isotope added while replicates refers to the number of samples within each condition and sums to 29 within each inoculant for a total of 87 across all three inoculants.

**Table S2.** Number of ASVs within each inoculant and strategy designation

| **Bacteria and archaea** | **Strategy designation** | **Isotope/ exudate treatment used for designation** | ***Betula*** | ***Eriophorum*** | **Permafrost** |
|---|---|---|---|---|---|
| | Exudate degrader | ASVs found in $^{13}$C exudates treatments but not in $^{18}$O-water with exudates treatments; ASVs found in both $^{13}$C exudate and $^{18}$O-water with exudates treatments | *306* | *29* | *189* |
| | SOM degrader | ASVs found in $^{18}$O-water with and $^{18}$O-water without exudates treatments but not present in $^{13}$C exudates treatments; ASVs found in $^{18}$O-water no exudates treatments but not $^{18}$O-water exudates treatments | 174 | 234 | 80 |
| | Exudate and SOM degrader | ASVs found in $^{18}$O-water with exudates treatments and $^{18}$O-water no exudates treatments | 55 | 15 | 7 |
| | Benefactor | ASVs found in $^{18}$O-water with exudates treatments but not $^{13}$C-labeled exudates or $^{18}$O-water no exudates treatments | 13 | 3 | 71 |
| | | Total non-unique ASVs across strategies | 548 | 281 | 347 |
| **Fungi** | Exudate degrader | ASVs found in $^{13}$C exudates treatments but not in $^{18}$O-water with exudates treatments; ASVs found in both $^{13}$C exudate and $^{18}$O-water with exudates treatments | 53 | 72 | 2 |
| | SOM degrader | ASVs found in $^{18}$O-water with and $^{18}$O-water without exudates treatments but not present in $^{13}$C exudates treatments; ASVs found in $^{18}$O-water no exudates treatments but not $^{18}$O-water exudates treatments | 21 | 51 | 3 |
| | Exudate and SOM degrader | ASVs found in $^{18}$O-water with exudates treatments and $^{18}$O-water no exudates treatments | 9 | 20 | 2 |
| | Benefactor | ASVs found in $^{18}$O-water with exudates treatments but not $^{13}$C-labeled exudates or $^{18}$O-water no exudates treatments | 0 | 0 | 0 |
| | | Total non-unique ASVs across strategies | 83 | 143 | 7 |

Strategy designation refers to the metabolic/ substrate preferences as revealed by comparing growth patterns across isotope and exudate treatments. *Betula, Eriophorum* and Permafrost refer to the inoculant community.

**Table S3.** Statistical tests comparing cumulative respiration and apparent priming effects across various inoculant and exudate treatments

| Group comparisons | F statistic | P-value | ω² effect size | Inoculant | Statistical test | Response variable |
|---|---|---|---|---|---|---|
| Exudate vs no exudate treatments | 23.9 | 1.21e-05 | 0.20 | *Betula*, *Eriophorum*, permafrost | Welch's ANOVA | Cumulative respiration |
| Exudate vs no exudate treatments | 0.09 | 0.77 | NS | Permafrost | Welch's t-test | Cumulative respiration |
| Exudate vs no exudate treatments | 95.8 | 6.19e-12 | 0.63 | *Betula*, and *Eriophorum* | Welch's t-test | Cumulative respiration |
| Exudate vs no exudate treatments | 6.34 | 0.021 | 0.03 | Permafrost | Welch's t-test | Respiration rate (day 46) |
| Inoculation vs no inoculation | 62.1 | 1.27e-10 | 0.45 | *Betula, Eriophorum,* permafrost | Welch's ANOVA | Cumulative respiration |
| Inoculation vs no inoculation | 81.8 | 4.79e-09 | 0.43 | *Betula, Eriophorum,* permafrost | Welch's ANOVA | Apparent priming |
| *Betula* vs *Eriophorum* | n/a | 0.98 | n/a | *Betula, Eriophum* | Games-Howel post-hoc | Cumulative respiration |
| *Betula* vs *Eriophorum* | n/a | 0.18 | n/a | *Betula, Eriophum* | Games-Howel post-hoc | Apparent priming |
| Inoculants | 76.5 | <0.001 | n/a | *Betula, Eriophorum,* permafrost | PERMANOVA | Total bacterial community composition |
| Inoculants | 1.20 | 0.289 | n/a | *Betula, Eriophorum,* permafrost | PERMANOVA | Total fungal community composition |

| | | | | | | |
|---|---|---|---|---|---|---|
| Inoculants | 1.28e2 | <0.001 | n/a | *Betula, Eriophorum,* permafrost | PERMANOVA | Active bacterial community composition |
| Inoculants | 1.60e3 | <0.001 | n/a | *Betula, Eriophorum,* permafrost | PERMANOVA | Active fungal community composition |
| Exudate vs no exudate treatments | 0.792 | 0.413 | n/a | *Betula, Eriophorum,* permafrost | PERMANOVA | Total bacterial community composition |
| Exudate vs no exudate treatments | 0.507 | 0.589 | n/a | *Betula, Eriophorum,* permafrost | PERMANOVA | Total fungal community composition |

Various statistical tests were conducted to test the differences in response variables across groups of interest. Group comparisons refer to the factors of interest for individual tests. F statistic represents the ratio of the variances between groups, p-value represents the probability of obtaining an F statistic due to chance and $\omega^2$ effect size represents effect sizes calculated using omega squared statistic. 'NS' designation refers to non-significant effect size whereas 'n/a' refers to not applicable. Inoculant refers to the source microbial community. Statistical tests included Welch's t-test, and ANOVA to compare the means across two or three groups respectively while accounting for unequal variances, Game-Howell post-hoc test for means separation, and PERMANOVA for non-parametric multivariate data. Response variables include cumulative respiration (µg C respired over the 54 day incubation), apparent priming (amount of C respired from exudate addition groups relative to controls), respiration rate (µg C * gram of dry soil$^{-1}$ * hour$^{-1}$) and microbial community composition.

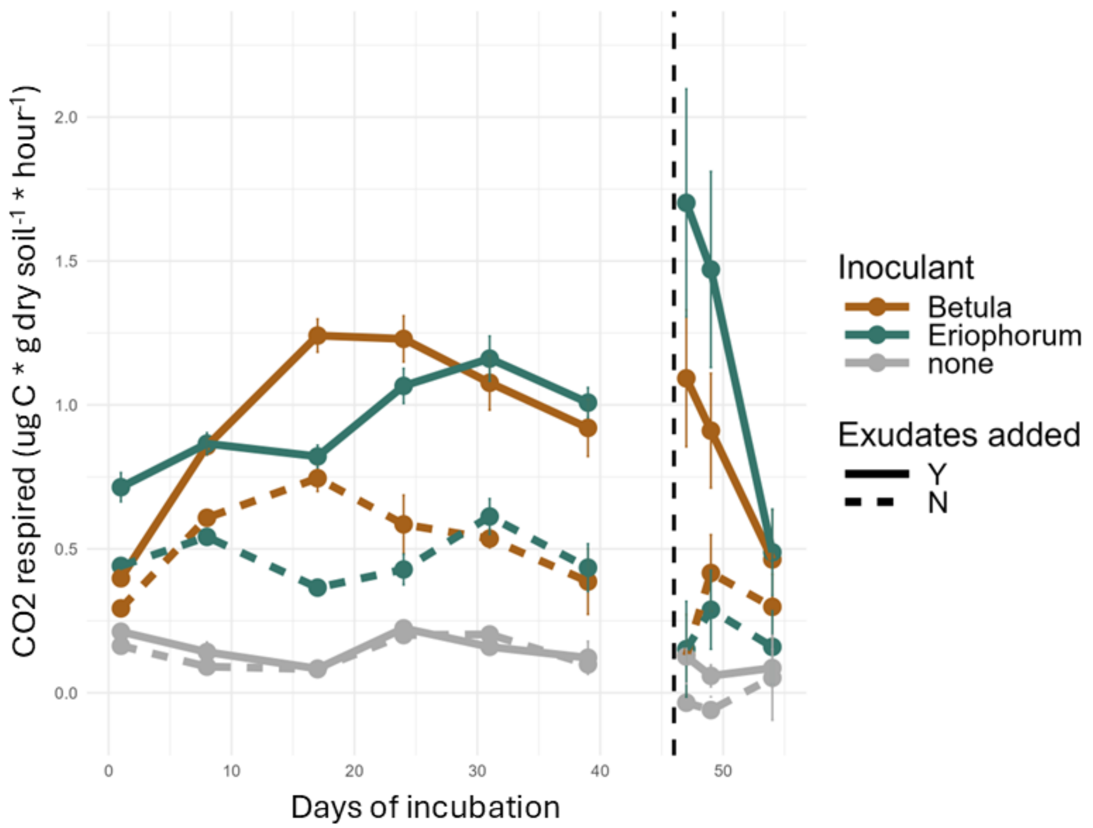


**Fig. S1.** Respiration rates were generally consistent throughout the experiment across daily additions (left panel) and after isotope spikes (right panel). *Eriophorum* (ERI) and *Betula* (BET) inoculants were the exception, as they increased their respiration rates immediately following the 7x C addition following the isotope spike before the resumption of their typical respiration rate. The respiration rate of uninoculated permafrost (PF) remained consistent with a minor, but statistically significant increase in respiration post isotope spike at day 46 ($F = 6.341$, p-value = 0.02148, Welch's ANOVA, $\omega^2$ effect size = 0.03).

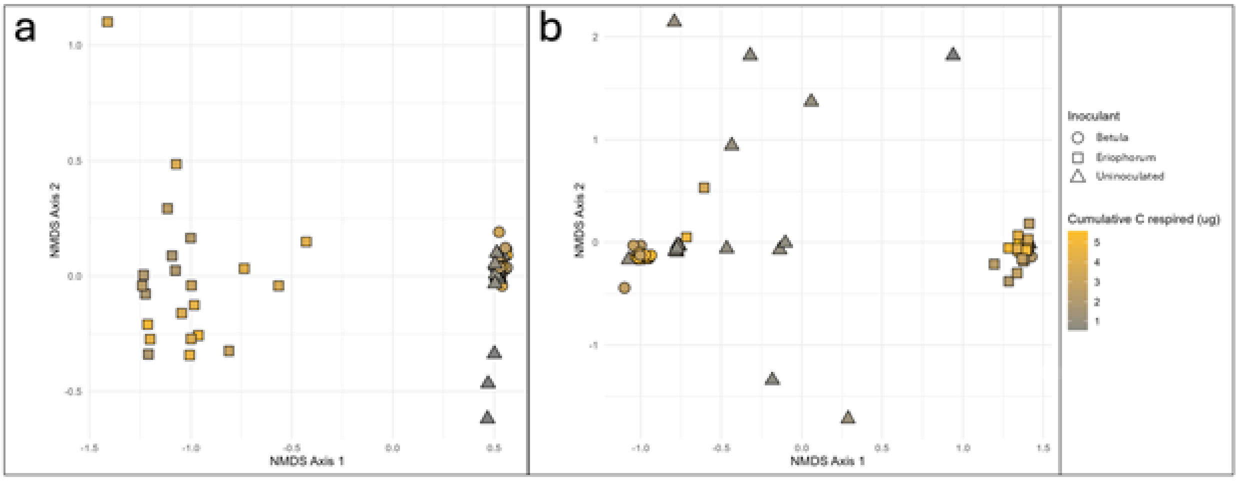


**Fig. S2.** NMDS of total microbial communities at the end of the incubation, which included active, dormant, and dead bacteria (a) and fungi (b) reveal distinct microbial communities associated with permafrost and rhizosphere inoculants. In total, *Betula* and *Eriophorum* respirations yield similar cumulative respiration values despite showing distinct community composition.

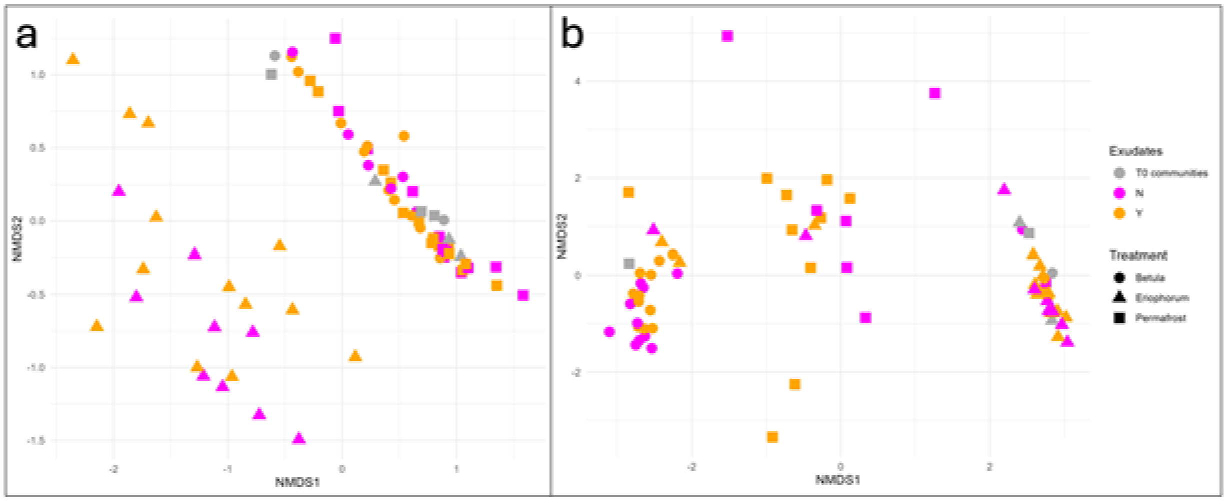


**Fig. S3.** NMDS of total microbial communities, which included active, dormant, and dead bacteria (a) and fungi (b) from the end of the incubation (Exudates N and Y), as well as initial community compositions after inoculation (T0 communities). The ordination reveals distinct patterns in microbial inoculant, however, does not show distinct patterns for exudate additions.

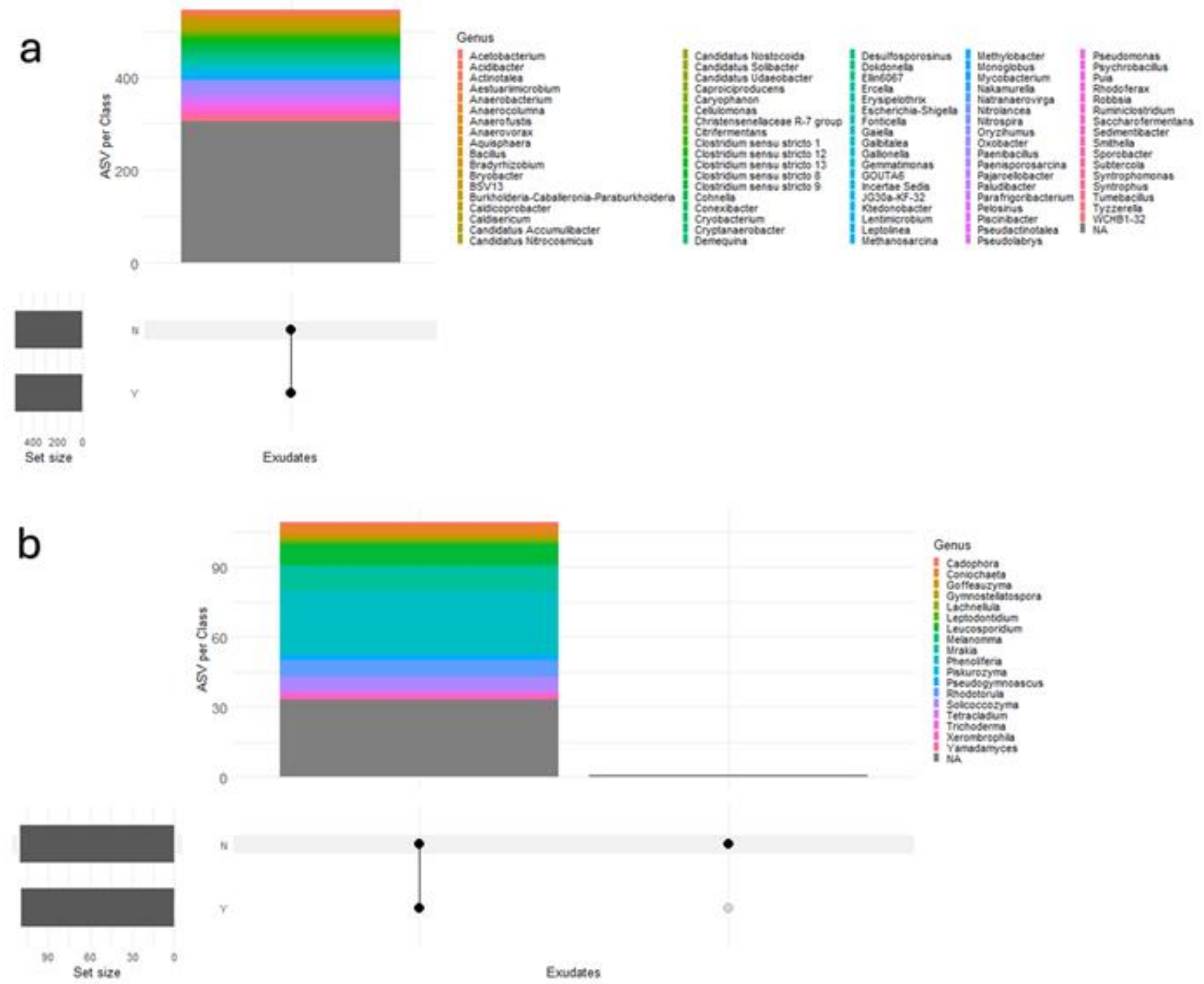


**Fig. S4.** Upset plots reveal that active bacteria (a) were shared across exudate addition and control groups, while few fungal taxa (b) were found in control groups not present in exudate addition treatments. To reduce the overinterpretation of the effects of rare taxa, we chose to only include taxa that contained 10 or more reads in our dataset. No exudates treatments contained one unique bacteria from the family Microbacteriaceae and two unique fungi belonging to the genera *Mrakia* and *Coniochaeta* that were filtered out of the Upset plot analyses due to low read counts.

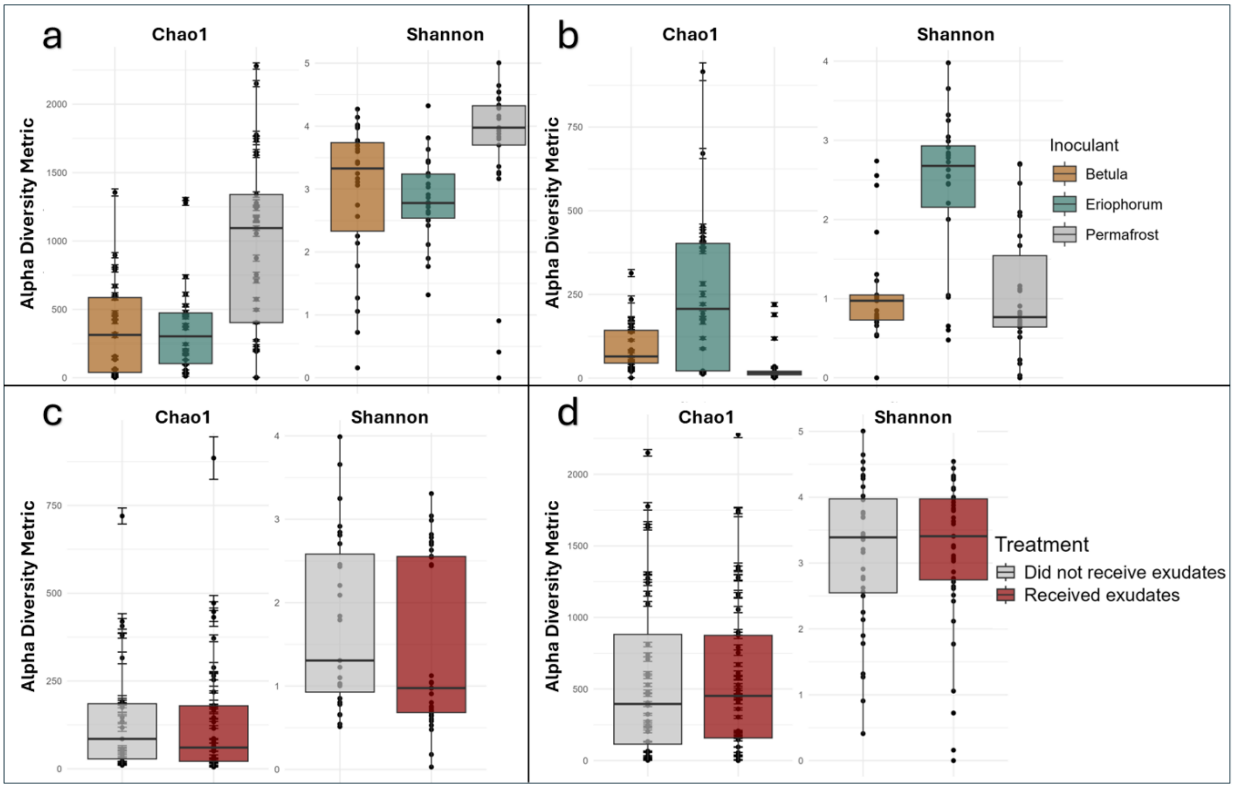


**Fig. S5.** Rhizosphere bacteria **(a)** and fungi **(b)** show distinct patterns of diversity based on the inoculants **(a, b)** but not exudate treatments **(c, d)**. Chao1 and Shannon represent diversity indices where Chao1 measures species richness and Shannon measures richness while accounting for species evenness.

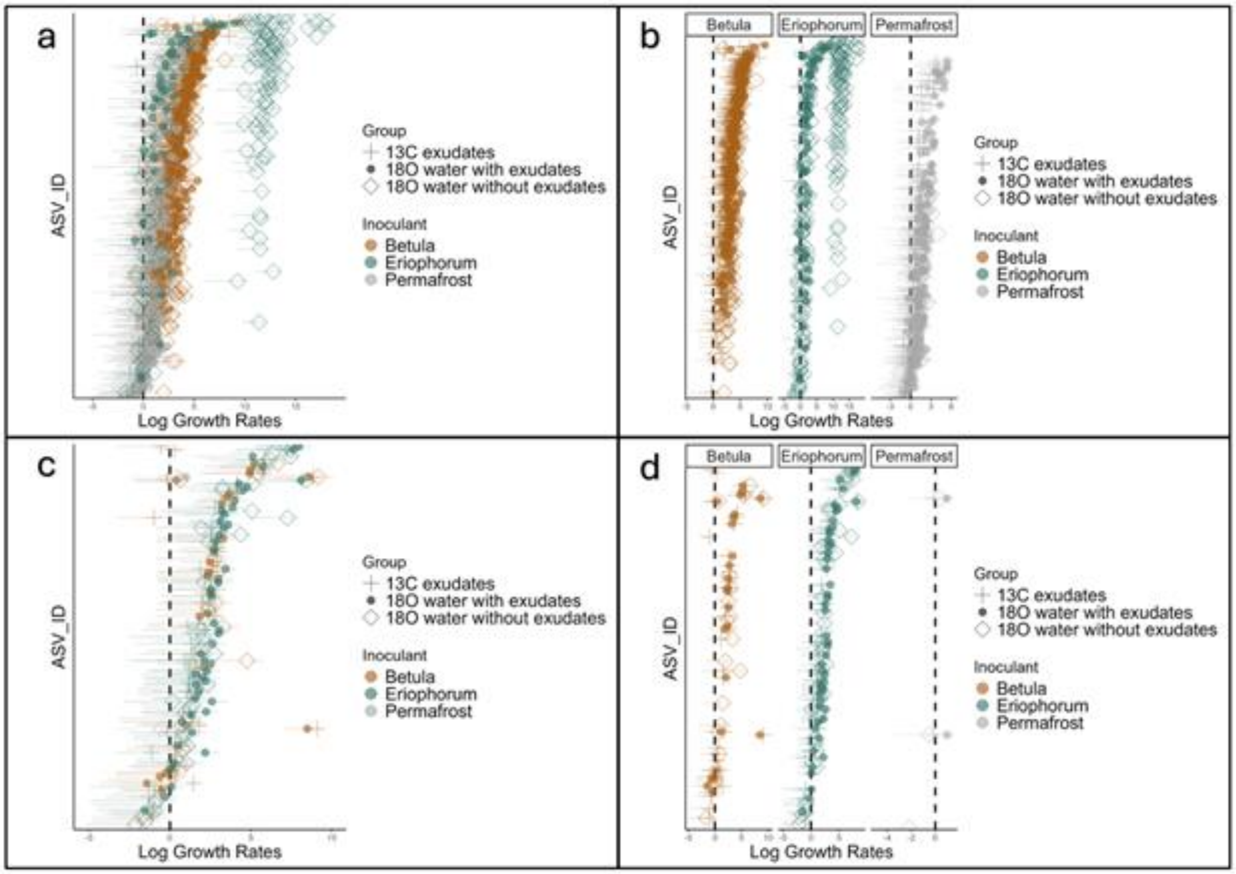


**Fig. S6.** Bacterial (a, b) and fungal (c, d) taxa highlight key differences in growth rates across isotope treatments (group) and rhizosphere inoculants. Each ASV is fixed along the Y-axis. Growth rates are displayed on the log scale and represent the number of gene copies per gram of dry soil per week. Columns representing *Betula*, *Eriophorum* and permafrost highlight the influence of community context on growth rates of individual taxa across communities. Confidence intervals on the tails of points represent the range of growth. Full plots (a, c) show the spread of taxon-specific growth rates while facets (b, d) show the spread of taxon-specific growth rates for each inoculant.

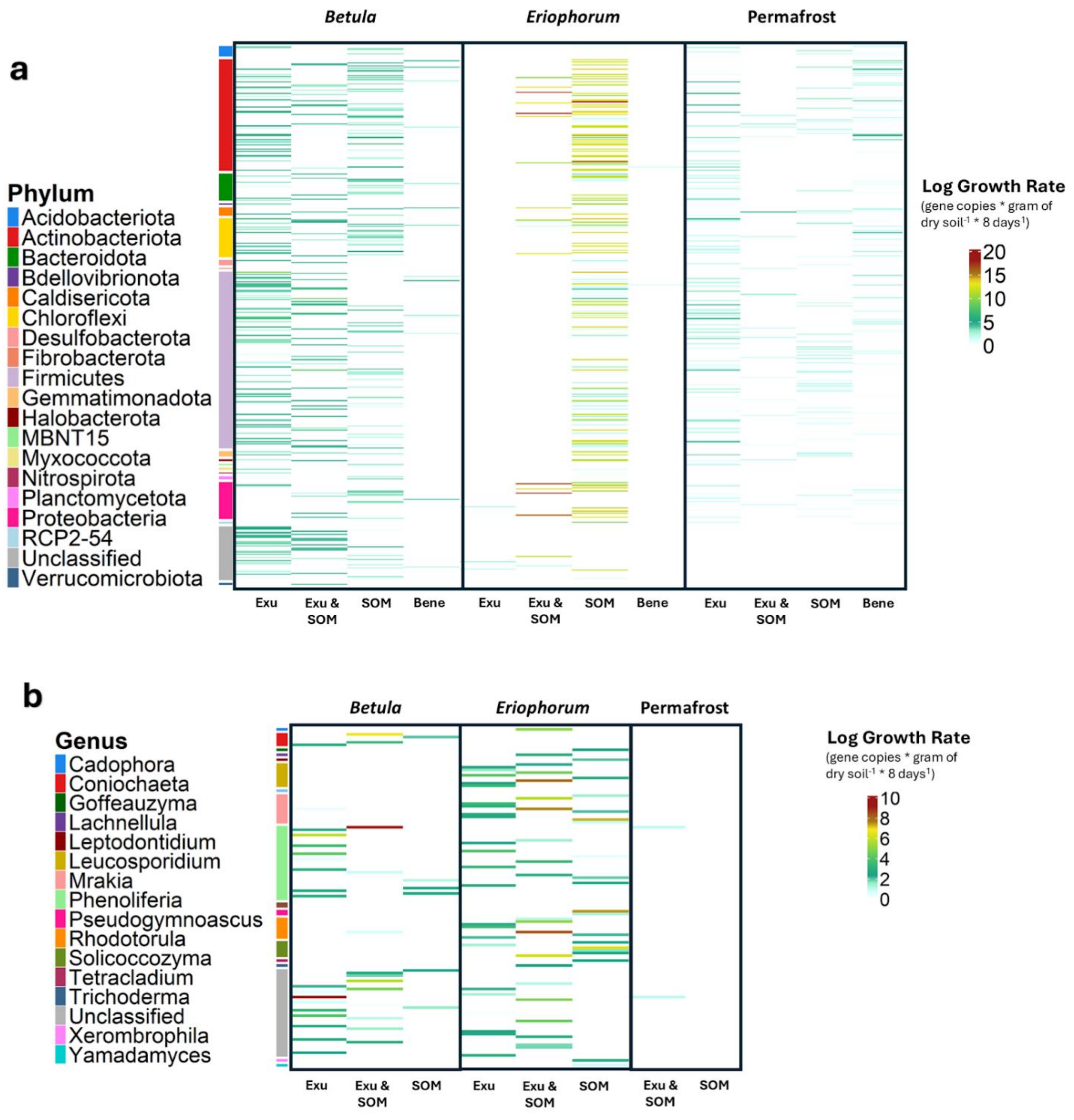


**Fig. S7.** Heat plots reveal distinct patterns of taxon-specific growth rates corresponding to inoculant communities and isotope treatments for bacteria **(a)** and fungi **(b)**. Each row is an individual taxon set in a fixed position across plots. Each column represents a microbial strategy: exudate degraders (Exu), exudate and SOM degraders (Exu & SOM), and SOM degraders (SOM). Columns are arranged by inoculants (*Betula*, *Eriophorum*, Permafrost). Growth rates are displayed on a log scale where warmer colors indicate fast growth and cooler indicate slower growth.

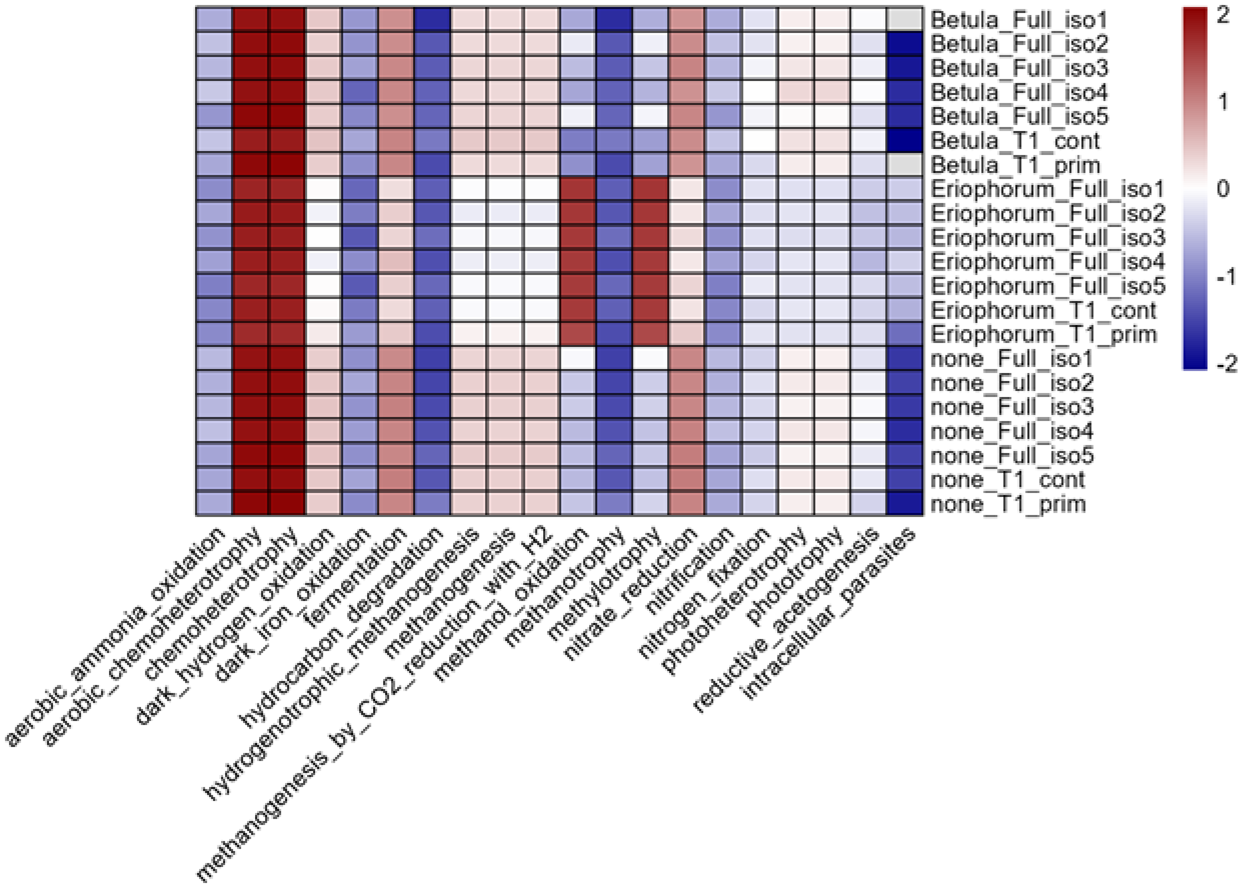


**Fig. S8.** Functional annotations of 16S sequences using FAPROTAX reveals consistent patterns across inoculants and exudate treatments. Iso numbers refer to the isotopic treatments where 1, 3 and 5 received exudates while 2 and 4 did not. T1_prim refers to pre-isotope treatments receiving exudates while T1_cont did not. Annotations were log scaled and relativized by each inoculant-isotopic grouping.
